# A membrane-driven biochemical oscillator tunable by the volume to surface area ratio

**DOI:** 10.1101/2025.11.14.688573

**Authors:** Jonathan Fischer, Ezra Greenberg, Yue Moon Ying, Sho S. Takeshita, Samuel L. Foley, Margaret E. Johnson

## Abstract

Oscillations are ubiquitous features of biological organisms, playing crucial roles in processes from circadian rhythms to developmental patterning. Protein-based biochemical oscillators have particular applications in synthetic biology because they can access fast and slow timescales that are independent from the transcription-translation machinery required of genetic oscillators. Here, we introduce and model such a mass-conserving biochemical oscillator using mass-action reaction kinetics that exploits dynamic changes to membrane phospholipid concentrations to drive proteins on and off the membrane in robust, tunable rhythms. Importantly, the oscillations rely on amplification of reactions on the membrane via dimensional reduction, and they are therefore tunable by variations in the volume-to-surface area ratio (V/A) of the system. With components inspired by the endocytic machinery, we show that a wide range of physiologically relevant biochemical rates can produce oscillations in part due to this independent geometric control. A broad computational screen of the high-dimensional parameter space reveals that oscillations require relatively strict enzyme kinetic design rules for low V/A but much more permissive kinetics for larger V/A. We validate that oscillations persist with more realistic reaction-diffusion simulations that captures explicit diffusion and stochastic, integer valued copy numbers, in overall good agreement with the period and amplitude of the deterministic oscillators. Because the oscillations rely on time-dependent changes to the surface properties and not post-translational modifications to the protein subunits, we demonstrate that it can be coupled to a self-assembling trimer, driving not only changes in localization but trimer yield. Our analysis establishes this membrane-localization oscillator as a new, geometry tunable and programmable timing module and suggests a potential for geometry sensing in engineered or cell-free systems.

## I. INTRODUCTION

Biological oscillators perform the crucial role of synchronizing biological processes—from circadian rhythms to cell cycle events—both internally and in response to external cues ^1 2–4^. Their ability to maintain robust temporal coordination even in the presence of significant molecular noise makes them highly attractive components for engineered biological circuits and synthetic applications. While genetic oscillators—such as the repressilator ^5^ traditionally dominated engineered oscillator designs, recent attention has shifted toward post-translational protein oscillators (PTOs)^6,7^. PTOs offer potential advantages such as faster dynamics (operating on millisecond-to-minute timescales compared to minutes-to-hours for genetic circuits), the ability to be targeted to specific subcellular locations, and potentially lower intrinsic noise ^8 9^. For instance, PTOs can be driven by rapid and reversible modifications like phosphorylation^10^, as seen in natural examples like the KaiABC circadian clock and eukaryotic clock cycles^11^, enabling operation on timescales independent of transcription–translation. Although promising models, such as a self-assembly-based post-translational oscillator ^12^ have been proposed, the overall toolbox for predictably engineering such dynamics remains limited compared to genetic circuits ^9^.

The dynamical principles underlying post-translational oscillator motifs are not limited to proteins; lipid species can similarly serve as substrates for modification states. Indeed, many cellular membranes exhibit dynamic interconversion among lipid species through phosphorylation and dephosphorylation. For example, the plasma membrane contains phosphatidylinositols (PIPs) that cycle among mono-, bis-, and tris-phosphorylated forms, with enzymes such as PI3K and PTEN regulating the interconversion between PI(4,5)P_2_ and PI(3,4,5)P_3_ ^13,14^. Similarly, in the Golgi apparatus, sphingolipids rapidly interconvert between ceramide, sphingomyelin, and glycosphingolipids. Membrane lipid dynamics are often critical in biological pathways dependent on phosphoinositides (PIPs)^15^. PIPs are essential for defining membrane identity (the “PIP code”) and for the selective recruitment of proteins to membranes, largely because their phosphorylation state determines binding specificity for various protein domains (e.g., PH, PX, C2) and influences electrostatic interactions. By modulating the local composition of PIP species, cells can finely tune membrane properties and control the recruitment of specific protein complexes. Moreover, enzymes regulating PIP phosphorylation— such as PIP5K and Synaptojanin^16^-- are themselves targeted to the membrane in a lipid-dependent manner, often involving specific protein interactions or lipid binding domains^17^. This targeted activity establishes nonlinear feedback loops between enzymatic action and PIP metabolism that can drive spatially propagating waves of proteins^15,18,19^. Importantly, localization to the membrane requires binding to the multi-component lipid bilayer, and this can therefore be a dynamic and tunable process. Protein domains—such as PH, PX, or ENTH domains—specifically recognize phosphorylated lipid headgroups^20,21^, meaning that shifts in membrane composition directly modulate protein binding. Consequently, by dynamically tuning the local lipid environment, cells achieve spatiotemporal control over membrane-protein affinities. This dynamic lipid landscape can encode complex molecular logic ^22,23^

Membrane localization can generate nonlinear feedback by selectively accelerating reaction rates and promoting assembly, due largely to the transition of diffusive searches from three-dimensional (3D) to two-dimensional (2D) spaces—a phenomenon known as dimensional reduction^24,25^. Dimensional reduction will enhance binding probabilities between binding partners localized in 2D vs if they remained in 3D when the volume-to-area lengthscale, *V*/*A*, is larger than the change in binding affinities between 3D and 2D, 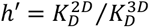. In other words, if we define a (dimensionless) dimensionality factor that combines these two changes, *DF*_*eq*_ = *V*/(*Ah*′), a *DF*_*eq*_ > 1 promotes more binding in 2D, with greater enhancement for larger *DF*_*eq*_. Although the kinetics of binding is more variable in being enhanced or impaired by 2D localization, both for receptor targeting^24^ and for self-assembly^27^, increases in concentration can typically overcome diffusive slowdowns on the membrane, supporting experimentally faster binding association in 2D^28^. The corresponding dimensionality factor for kinetics is thus dependent on the ratio of 3D and 2D rates, 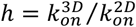. With 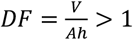, association reactions between species localized to 2D will be accelerated relative to their counterparts in 3D.

Despite extensive study of lipid–protein interactions, the application of the lipid membrane’s potential for geometrically tunable, nonlinear feedback in driving *synthetic* biochemical oscillations has remained largely unexplored. We term this potential class membrane-localization oscillators (MemLOs). To address this gap, our study develops a minimal model for a biochemical oscillator driven by membrane localization—one that leverages the geometry of the reaction system itself relative to the 2D rates (*DF*) to coordinate reaction dynamics. Our design draws inspiration from the PIP-mediated process of clathrin-mediated endocytosis (CME). In CME, the phosphorylated lipid PI(4,5)P_2_ functions as a key signal, recruiting the central adaptor protein AP2^29^. While endocytosis is not an oscillatory system, the frequency and stability of endocytic events is clearly dependent on the composition of the membrane, both PIP2 and receptor density^30^. Phospholipid kinases and phosphatases are both known to bind directly to the central adaptor protein AP2^31^, and while phosphatases are required for later trafficking steps^30^, these lipid enzymes could also help to locally tune the stability of clathrin assemblies on the membrane.

To characterize the dynamical properties of our oscillator and assess its experimental feasibility, our study is organized as follows. We first determined the regions of parameter space that can produce oscillations, using genetic algorithms to widely explore biologically feasible rate constants, initial monomer concentrations, and dimensionality factors (DF, representing the V/A ratio) for parameters that can produce oscillations. Next, we examined the tunability of the oscillation period and amplitude and analyzed the robustness of these traits to noise and perturbations to the parameter values. We then compare nonspatial modeling outcomes with a spatial reaction–diffusion simulation and demonstrate how protein oscillations in our system can be coupled to higher-order self-assembly. Finally, we discuss the implications of our findings for experimental synthetic biology.

## II. MODEL and METHODS

### II.1 Model Description

Our model contains 5 monomeric species, an adaptor protein *A*, a phosphatase *P*, a kinase *K*, and a lipid that can exist in two states, unphosphorylated *L* (e.g. PI(4)P), and ‘sticky’ phosphorylated lipid *Lp* (e.g. PI(4,5)P_2_). They react with one another via only bimolecular and unimolecular reactions, with 5 distinct reversible binding reactions and two distinct (irreversible) catalytic reactions, necessitating 5 on-rates 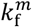, 5 off-rates 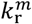, where *m* indicates which reactant monomers participate, and 2 catalysis rates 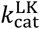 and 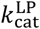 for enzymes *K*, and *P* (Fig 1). The proteins The enzymes K and P each have two binding interfaces, one for the lipid substrate (L and Lp respectively), and the other to bind to the adaptor protein A. Similarly, A has two binding interfaces, one to bind to the membrane via Lp and the other to bind an enzyme. Note that A can only bind one enzyme at a time. The lipids each only have a single interface for binding proteins, but they can be catalytically converted to the other state by the appropriate enzyme. In our microkinetic model, we **allow** all possible intermediates, and can thus form complexes containing 2-4 monomer species, producing 16 distinct species total (Fig 1c). With 4 mass conservation equations for the three proteins and lipid (L+Lp), there are thus 12 independent variables. The full reaction network contains 19 reversible reactions and 6 catalysis reactions. However, aside from dimensional reduction, we assume no added cooperativity in the reactions, **so** the same rates are used for the same interface interactions between higher-order intermediates. For analytical **treatment** of our reaction network, we also defined a restricted model which removed some cytosolic interactions to reduce our system to 12 distinct species and thus 8 independent variables (see SI).

**Figure 1.**
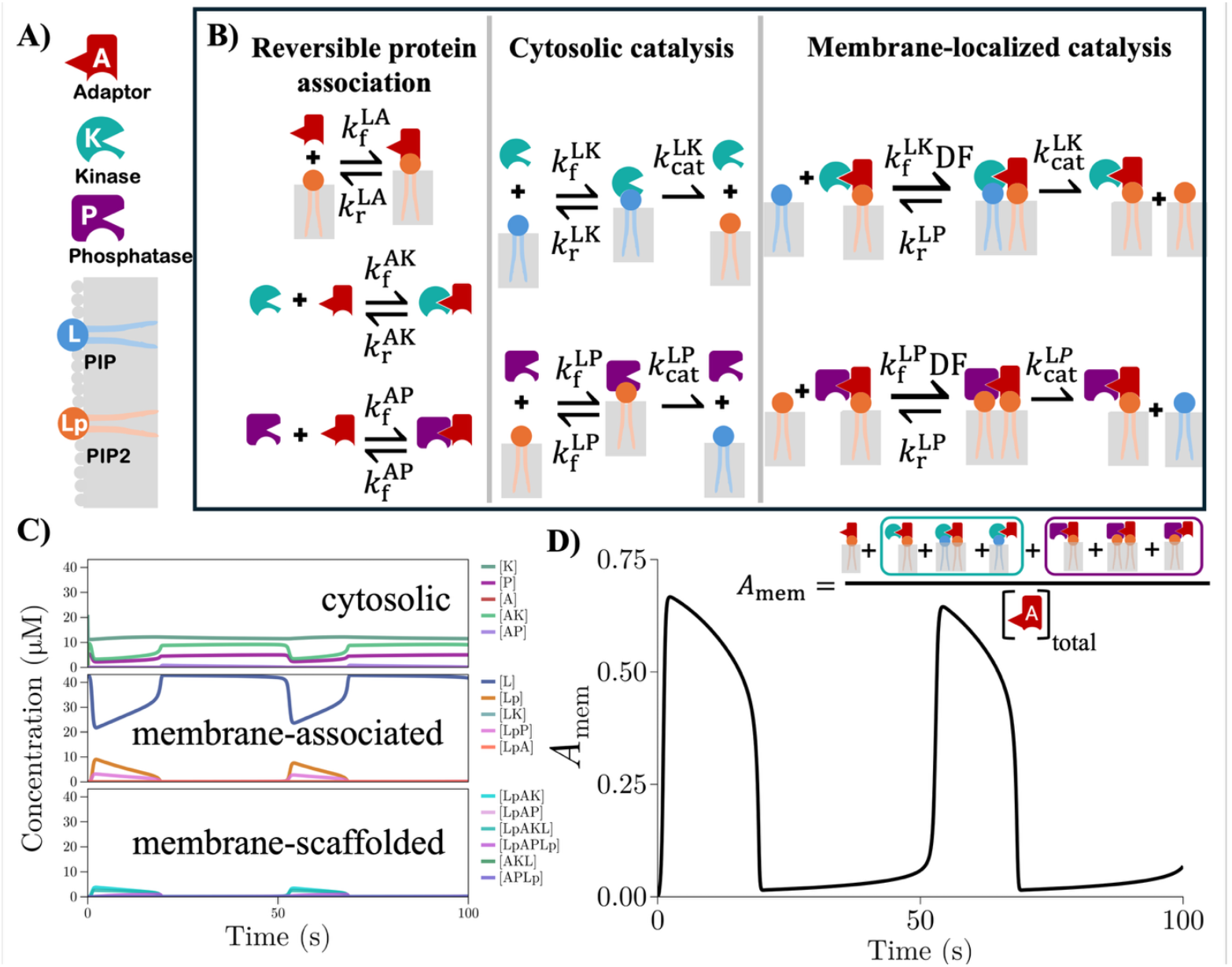
Model for the membrane-protein oscillator driven by lipid-modifying enzymes. A) The model contains 3 protein species, the adaptor (red) and two lipid enzymes (green and purple). Each protein has two interfaces, allowing for complex formation with up to 4 species. The lipid specie has one interface and is restricted to the 2D membrane but in two states, the inactive L (blue) and the ‘sticky’ phosphorylated Lp (orange). B) The reaction network builds from the key fundamental reactions shown in the first two boxes of 5 reversible reactions and 2 irreversible catalytic reactions, and thus has 12 tunable rate constants plus the dimensionality factor DF. The adaptor only binds Lp and not L, creating asymmetry. The third box illustrates how the enzyme association reactions transform to 2D reactions, with rates scaled by DF. This requires co-localization of the enzymes by the adaptor bound to the sticky lipid Lp. C) An oscillatory solution showing all 16 species concentrations vs time, classified based on whether they are purely cytosolic (top), lipids or protein-bound lipids (middle), or higher-order 3-4 component species (bottom). D) Our primary observable (*A*_mem_, black) would mimic an experimentally tagged adaptor protein. *A*_mem_ reports the fraction of adaptor on the membrane relative to the total concentration of adaptor.

We model the dynamics of our system using mass-action kinetics and numerically solve the resulting systems of ordinary differential equations (ODEs) in Julia with the Rodas5P integrator. To explicitly assess the roles of diffusion, spatial inhomogeneities, and stochasticity, we also evaluate multiple solutions using stochastic, particle-based reaction diffusion simulations propagated with the NERDSS software^32^. The ODEs were integrated to 2000 seconds, which we chose to provide a reasonable balance of computational efficiency and orders-of-magnitude variation in oscillation periods. We note that longer periods are feasible given slow rates (<10^-3^s^-1^) and low concentrations (<0.1uM) but would not detected. While our reaction system includes bindingin solution (3D),to membranes (3D-to-2D), and restricted to the membrane (2D) (Fig 1),for our ODE models we track all species as bulk (3D) concentrations with typical V^-1^ units of μ*M* by accounting for dimensional reduction, or the transformation of 3D interactions to a 2D search space.This changes both the effective i) concentrations and ii) rate constants. For (i), bimolecular association in 2D involves a search between species restricted to the membrane area *A* with densities of, e.g. *ρ*_*X*_ = *N*_*X*_/*A*, with *N*_*X*_ copy numbers. In solution, the concentration is [*X*] = *N*_*X*_/*V*, and therefore they are related via 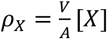, meaning 2D densities are captured by rescaling relative to a bulk concentration by 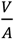. For (ii), 2D association reactions are parameterized by 2D rates constants (although in 2D they are only approximately constant^33^), which are most compactly related to their 3D counterparts by a lengthscale 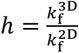 ^34^, and have been measured in the 1-100 nanometer range^28,35^. The magnitude of *h* depends on the specific binding pair. The off-rates can also change but only by a dimensionless scalar *c*, such that 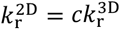. We will assume the off-rates are unchanged to minimize additional parameters (*c* ≡ 1), and thus we have 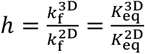. This now thermodynamically expressed lengthscale has been more extensively studied, and in the simplest approximation is entirely defined by entropic effects due to loss of rigid-body degrees of freedom, putting it again in a molecular or nanometer scale^36–38^. However, for proteins that are enthalpically favored to interact in 2D, the lengthscale can be orders-of-magnitude smaller, and thus it does provide tunability, albeit coupled to allosteric changes on the surface^39^. Collectively we combine these effects into a single dimensionality factor,

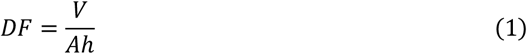

To minimize additional parameters, we also assume the same value for all 2D reactions (SI for further details), although it can vary between pairwise species subject to constraints due to thermodynamic cycles in our model (see SI). Since we assume the same off-rates in 3D and 2D, it is therefore the forward association reactions that must be scaled, such that 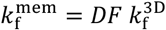 given 3D concentrations, and this has units of μ*M*^−1^*s*^−1^. If we assume that the V/A ratio is within physiologic values expected for cells (∼ R/3 or ∼0.5-10 μm)^40^, and then consider values of h between 1nm to 500nm, we can sample DF values from 1-10000, which is the range we consider here. For spatial simulations, *h, V*, and *A* must each be separately specified, which can impact results, primarily due to diffusion times to the membrane surface for large V/A.

#### Trimer extended model

For the coupled heterotrimer self-assembly model seen in Figure 6, we defined two additional ratiometric observables. *Trimer Yield* measures the overall efficiency of the assembly process, defined as the total concentration of assembled trimers divided by the initial concentration of the limiting monomer *M*_lim,0_:

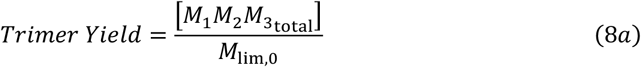

where 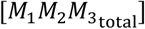 is the sum of all cytosolic and membrane-bound trimer species. *T*_mem_ measures the spatial control exerted by the oscillator, defined as the fraction of fully assembled trimers that are localized to the membrane:

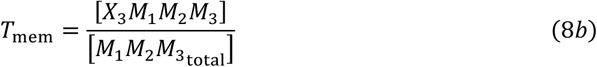

where [*X*_3_*M*_1_*M*_2_*M*_3_] is the fully membrane-anchored trimer complex.

### II.2 Phenotype measurements

We measure the ‘phenotype’ or period and amplitude of a solution from the observable *A*_*mem*_(*t*) ∈ [0,1], which tracks adaptor species localized to the membrane relative to the total adaptor species, facilitating comparison between models with varying initial concentrations and providing a useful metric to track experimentally, given a ‘tagged’ adaptor. For the restricted model:

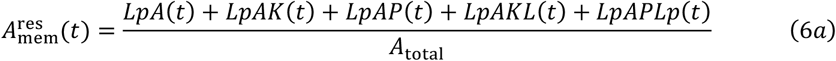

For the full model:

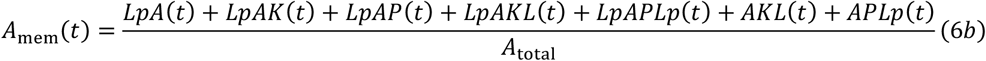

The period and amplitude of *A*_mem_ are computed with the following expressions:

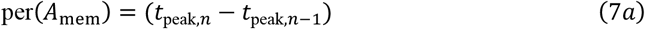

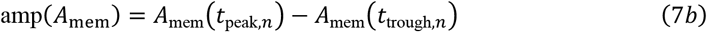

where n refers to the index of the last peak. We exclude the first 10% of the time points when evaluating the oscillatory behavior as it is impacted by the initial conditions relative to subsequent periodicity as the initial trajectory is highly dependent on the initial conditions and often does not reflect the limiting behavior of the system. Further, we found that this helped to prevent the genetic algorithm from propagating solutions with damped oscillations.

### II.3 Optimization with a genetic algorithm

For reasons of evolvability, the restricted observable variant 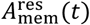, which omits AKL and APLp from the numerator, was computed in addition to the full variant and used solely for producing and scoring the actual FFT spectra during fitness function evaluation. Doing this improved the smoothness of the fitness landscape during early generations of the genetic algorithm, and thus the diversity and abundance of oscillatory solutions found. All other solution phenotype properties computed both during fitness evaluation, like period and amplitude, or in downstream analysis, used the full and complete *A*_mem_ solution vector.

To generate solutions with stable oscillations, we optimized the model inputs, or genotypes, using a genetic algorithm (GA) implemented in the Julia package Evolutionary.jl. By operating on a population of randomly generated initial genotypes and selecting for ‘fit’ individuals over successive generations of mutation, crossover, and selection, GAs can naturally produce a broad set of fit solutions, rather than local optimization techniques that converge to a single minima. Initial values for the 17 parameters of each individual are selected based on a log10 uniform distribution within physiologic limits for rates (SI Table), with concentrations restricted to 0.01-10uM. We found that the most important hyperparameters of the GA optimization were the population size and the number of generations, which for tractability were set to 100,000 initial random individuals evolved for 5 generations, producing 600,000 total solutions. Other hyperparameters were systematically tested but showed less impact on the overall size of the solution space found (see SI).

#### Fitness function

The fitness function for assigning scores to oscillatory behavior used properties of the real-time ODE solution 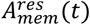 and its discrete Fourier transform, 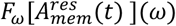. We found empirically that for lower DF, we found larger solution sets using 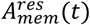, but for the analysis of the oscillator amplitudes, we report values from the full species *A*_*mem*_(*t*).

For all solutions 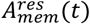, we evaluate their discrete Fourier transform 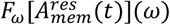 using the built-in Julia function rfft. We calculate the one-sided complex modulus of the DFT. To speed up the computation, we use slightly fewer data points (N=19,683) than the 20,000 point real-space vector, and normalize the resulting DFT array by half its length (N/2). Given the numerical complex modulus, we use the built-in finite difference peak finder algorithm (using default parameters) to identify *N*_*P*_ peaks in the spectrum. For each *i* peak we evaluate its amplitude *γ*_*i*_, and its sharpness s_*i*_. The value s_*i*_ reports the standard deviation of the peak height at its maximal amplitude compared to the amplitude at the neighboring point on either side. We ignore peaks with a height <1E-2. A larger standard deviation indicates a more distinct peak, suggesting a stronger and more stable oscillatory behavior. We then calculate the fitness, we are trying to maximize, using

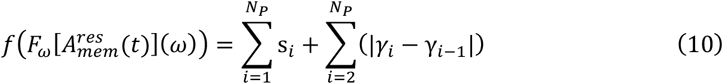

The second term selects for high-amplitude oscillations at a single frequency. This function identifies short and long period oscillators, with a range of amplitudes. High fitness is highly predictive of undamped oscillators. These fitness values are used by the genetic algorithm to evaluate which individuals to breed, propagate, mutate.

#### Classifying Solutions as Oscillatory for Further Analysis

After running the genetic algorithm, we use a series of heuristics to classify each ODE solution as either oscillatory or not, and we typically save only the oscillatory solutions for further analysis. These heuristics are based on the actual trajectories of the A(t) concentrations rather than the spectra in Fourier space. First, a first-order difference peak finder algorithm was applied to the 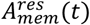 trajectories, and any solution with less than two peaks (or local maxima?) in the 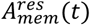 trajectory is discarded. As stated earlier, this limits the maximal period of oscillations that we can detect to <2000s. Next, we screen out solutions that show very low magnitude amplitude: if the amplitude of oscillations in 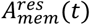 is less than 0.01, corresponding to 1% of the total concentration of 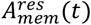, we also treat the solutions as non-oscillatory. We then apply a third screen to discard solutions with damped oscillations. If the last 200 s of 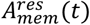 have a standard deviation <1e-6 μM, indicating the A(t) converges to a steady state, the solution is classified as non-oscillatory.

For the remaining candidates for oscillations, we perform a final screen to ensure that the solutions show regular periodicity and amplitudes. First, we apply a prominence-based peak detection algorithm (with a minimum prominence threshold of 0.01) to the time series of *A*_*mem*_ to identify the indices of local peaks and troughs. Next, we assess the regularity of the oscillation by comparing both the peak amplitudes and the temporal distances from each peak to its nearest trough. Let (*h*_*i*_) denote the amplitude of the *i*th peak, and let *d*_*i*_ denote the time interval (peak-to-trough distance) associated with that peak. We compute the fractional variation in amplitude between two consecutive peaks as 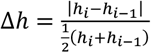 and similarly, the fractional variation in the peak-to-trough distance as 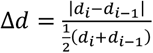. A candidate solution is accepted as oscillatory only if there exists at least one pair of consecutive peaks for which both Δ*h* and Δ*d* are less than 0.01 (i.e., a 1% variation threshold). This criterion—checked post-hoc on the optimized dataset on every candidate individual—ensures that the oscillation exhibits both consistent amplitude and regular timing, thereby filtering out spurious solutions that might otherwise pass the initial criteria, especially in regimes with low dimensionality factors.

### II.4 NERDSS simulations

To understand the impact of spatial variation and discrete molecular copy numbers, we performed structure-resolved reaction-diffusion simulations using the Non-Equilibrium Reaction-Diffusion Self-assembly Simulator (NERDSS) software package. NERDSS generates rigorous stochastic assembly trajectories using the Free-Propagator Reweighting algorithm. Individual molecules are represented as center of mass sites with associated coarse-grained binding sites. The reaction network is identical in NERDSS to that of the ODE system and is parametrized by microscopic on- and off-rates (*k*_*a*_, *k*_*b*_), which are calculated from macroscopic rates (*k*_*on*_, *k*_*off*_) by incorporating molecule diffusion constants, which we provide with the molecule geometry (see SI). The volume *V* and surface area *A* are explicitly represented in NERDSS simulations via a rectangular prism simulation cell with specified dimensions. 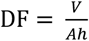 is then determined by the addition of the parameter *h*, which relates a 2D (microscopic) bimolecular reaction rate to its 3D counterpart in NERDSS, 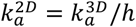. For simplicity, we take *h* to be 1nm throughout unless specified otherwise. The system volume *V* is chosen so that there are at least 100 copies of the lowest concentration species. With these conventions, each ODE parameter set uniquely determines a NERDSS simulation up to choice of timestep, for which we use *δt*=10μs unless specified otherwise. We tested 25 ODE parameter sets with oscillatory solutions; those which successfully exhibited oscillations in NERDSS replicated with 5 independent runs.

#### Period and amplitude measurement

To robustly quantify the period and amplitude of the stochastic trajectories from NERDSS (Fig. 4B, C), we used alternative methods for measuring period and amplitude from those used against deterministic ODE solution.

The oscillation period for each stochastic simulation was determined from its power spectrum, which quantifies the dominant frequencies in the *A*_mem_(*t*) signal. A three-step filtering process was then applied to the spectrum to robustly identify the fundamental frequency and resulting period. First, noise was filtered out, defined as frequencies less 25% of spectrum’s maximum power. Next, the remaining frequencies was separated from their subharmonics. This harmonic filtering was performed by testing each candidate frequency individually. A candidate was rejected as a subharmonic artifact if its power was found to be less than the power of its own first harmonic, located at twice its frequency. From the set of candidate frequencies that passed both the noise and harmonic filters, the one with the highest power was selected as the fundamental frequency. The period was then calculated as the inverse of this frequency. If no candidate passed the full filtering process, the period was derived from the single frequency with the maximum power in the spectrum.

A similarly robust method was used to measure the oscillation amplitude. To avoid sensitivity to stochastic noise spikes, the peak-to-trough amplitude was defined as the difference between the 98^th^ percentile and the 2^nd^ percentile of the entire *A*_mem_(*t*) time-series. This percentile-based approach captures the core range of the oscillation while effectively disregarding the most extreme 2% of data points at the top and bottom of the range as outliers.

### II.5 Predicting Period with a Multilayer Perceptron

To predict period from the input parameters a multilayer perceptron consisting of 5 layers of 128,128,18,128, and 128 neurons respectively was constructed. The middle layer utilizes an Information Ordered Bottleneck layer{Ho, 2025 #2507} in which during training all neurons are hierarchically unmasked. All the layers except the middle layer and last layer are followed by a leaky ReLU activation. All weights were initialized using a Kaiming normal distribution and the bias for the output was initialized to the mean period of the training set.

The data was first stratified with DF with a 70/30 split and then the 30 split was randomly split 2/1 for a train/validation/test split of 70/10/20. The 17 input parameters (k_f_^LA^,k_r_^LA^,k_f_^LK^,k_r_^LK^,k_cat_^LK^,k_f_^LP^,k_r_^LP^,k_cat_^LP^,k_f_^AK^,k_r_^AK^,k_f_^AP^,k_r_^AP^,DF,L,K,P,A) were processed via a log transformation and then Z-score normalized. The Period was log transformed and then min-max normalized.

For hyperparameters, a batch size of 1024, 1000 epochs, a learning rate of .001, a Mean Square Error (MSE) loss, and the AdamW optimizer were used. Training was done with 4 A100 GPUs and early stopping after no improvement in validation loss for 50 epochs. Validation loss was evaluated every 10 epochs.

### II.6 Symbolic Regression for Model Discovery

To identify an interpretable mathematical model describing the relationship between the 17 input parameters and the period, we employed symbolic regression (SR). This analysis was conducted using the PySR (Python SR) library{Cranmer, 2023 #2508}, a tool that utilizes an evolutionary algorithm to search the space of mathematical expressions to find the optimal equation to model data.

A PySR regressor model was configured with a specific set of operators, hyperparameters, and constraints to guide the evolutionary search. The search algorithm was permitted to construct candidate equations using a predefined set of mathematical functions. The allowed binary operators were addition, subtraction, multiplication, division, and exponentiation. The allowed unary operators were cosine, natural exponential, sine, inverse, natural logarithm, absolute value, square root, square, cube, base-10 logarithm, and cube root.

The fitness of candidate equations was evaluated by minimizing the MSE. The evolutionary search was configured to run for a total of 1000 iterations, with 500 population cycles performed within each iteration. To control the complexity of the resulting expressions, the maximum number of nodes (operators, variables, or constants) in any single equation was limited to 30, and the maximum nesting depth of operators was limited to 5. Early stopping was used to halt the search if an equation was discovered with a MSE loss of less than 10^−6^ and a complexity score of less than 10. Training was done on the validation dataset from the period predictor training.

To prevent redundant or non-physical equation structures, several constraints were defined. A nested constraint was defined where the operator exponent of *e* was prohibited from being a direct child argument to the square, cube, or exp operators. This explicitly prevents the formation of expressions such as (*e*^*x*^)^2^, (*e*^*x*^)^3^, and 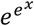, which can be simplified or lead to unnecessarily complex structures. A power operator constraint was also applied, where the operator was constrained to expressions of the form (*constant*)^(*variable*)^. Expressions of the form (*variable*)^(*constant*)^ are already handled by the square and cube operators or can be evolved through multiplication. The following expression had the lowest loss and was used to plot Fig. 4: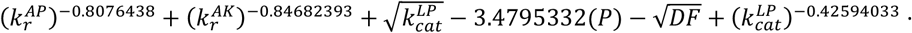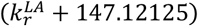

## III. RESULTS

### III.1 Adaptor proteins oscillate on and off a membrane with a dynamic lipid composition

The adaptor protein A in our model binds selectively to only one of the two lipid states, Lp, mimicking the behavior of diverse lipid-binding domains^21^. Therefore, creating time-dependent oscillations in the Lp species can produce cycles of adsorption and desorption of A to and from the membrane, which we label *A*_mem_(*t*) as it comprises multiple microstates (Fig 1). A key feature of our model is that the oscillations in the lipid species are themselves dependent on recruitment by the adaptor protein, such that only three distinct protein types are present in the model (A, K, P), in addition to the membrane-restricted lipids (Fig 1). Oscillations emerge in our model primarily from two mechanisms. First, feedback is generated by the membrane-localized adaptor, LpA: it localizes the enzymes to the membrane where their activity is amplified by the 2D search for their lipid substrate. The phosphatase P produces negative feedback by removing Lp from the surface and thus driving A to desorb and take the enzymes with it back to 3D. The kinase K generates positive feedback by creating Lp and thus enhancing adaptor (and thus enzyme) adsorption. Second, the negative feedback is delayed because it is mediated by the multi-step formation of the adaptor-phosphatase intermediate. This delay is essential for sustained oscillations ^1^. The adaptor competes with the membrane-localized phosphatase for the substrate Lp and the adaptor cannot bind phosphatase when K is bound (Fig 1).

### III.2 Adaptor protein and both enzymes are necessary for oscillatory lipid behavior

Our model suggests two fundamental requirements for sustained oscillations: the membrane-localizing adaptor protein to structure the positive and negative feedback loops, and a sufficient amplification (quantified by DF>1) of reaction rates in 2D, which we return to below. When the adaptor is removed entirely, our model reduces to a well-studied cycle of post-translational conversion of substrates (L, Lp) by their counteracting enzymes (K, P). This system does not produce oscillations in any species ^2^, following analysis of chemical reaction networks (CRN: this assumes the system is well-mixed) ^41,42^. If the adaptor is added to this system to reversibly bind phospholipid Lp but without interacting with either enzyme, the stability of the steady-state behavior does not change (Table 1, see SI). Thus, the production of oscillations requires an adaptor to bind to at least one of the enzymes. Further, because our system is mass conserving, both enzymes must be present in the network to allow the lipids to cycle repeatedly between both L and Lp states, otherwise the lipids will be irreversibly converted to a single type.

**Table 1.** (QSS) means the results are specific to our reduced, approximate model. **\***This requires that we enforce (as we do here) that binding to the adaptor cannot change the enzyme activity through some alternate allosteric mechanism. Then it reduces to the case of A_0_=0.

| MODEL | Ablations | Analytical analysis:<br>oscillations? | Numerical<br>sampling:<br>oscillations? |
| --- | --- | --- | --- |
| Full model | - | - | Yes |
| **Restricted model** | AK and AP only<br>form on membrane | - | Yes |
| **Two-variable**<br>**Reduced model** | Quasi-SS<br>approximations | Yes | Yes |
| **No adaptor-enzyme**<br>**interactions** | $k_f^{AP} = 0 \text{ \& } k_f^{AK} = 0$ | No | - |
| **No adaptor-**<br>**phosphatase binding** | $k_f^{AP} = 0 \text{ \& } [LpAP]=0$ | No evidence (QSS) | - |
| **No adaptor-lipid**<br>**binding** | $k_f^{LA} = 0$ | No* | No |
| **No membrane-**<br>**driven enhancement**<br>**(DF = 1)** | $k_f^{LK,2D} = k_f^{LK}$<br>$k_f^{LP,2D} = k_f^{LP}$ | No evidence (QSS) | No |

### III.3 Enzyme reactions must be ‘activated’ by localization from the adaptor protein

While both the enzymes and the adaptor protein are necessary for lipid oscillations to be possible, the mere presence of all these proteins appears insufficient to generate oscillations: the enhancement of enzyme activity by membrane localization is also necessary. Mathematically, we find evidence that the DF must be greater than one, indicating faster reactions on the membrane, for oscillations to occur. Analysis of the graphical structure of the CRN does not help in establishing oscillations in this case (SI), so we performed a reduction of our restricted, 8-variable reaction network using quasi-steady-state approximations (see SI for full derivation). By systematically applying quasi-steady state (QSS) assumptions to fast intermediate species, the model complexity was reduced to a two-variable ordinary differential equation (ODE) system describing the dynamics of the dimensionless lipid concentrations 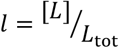 and 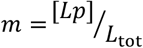. This reduced two-variable system, despite involving mathematically complex rate functions, qualitatively recapitulates the oscillatory behavior of the full and restricted models under specific parameter regimes, confirming that the reduction captured some essential dynamical features. The primary utility of the two-variable system lies in its amenability to phase plane analysis techniques that can rule out oscillations under specific conditions. We show by induction that a further simplified 2-variable model does produce oscillations in specific parameter regimes, with DF=80 for example (SI). We performed linear stability analysis by examining the Jacobian matrix of the reduced *l, m* system and applied the Bendixson-Dulac criterion (specifically, Corollary 3, which examines the sign of the Jacobian trace). First, by setting the dimensionality factor to unity (DF=1), we observed that the trace of the Jacobian is negative in the valid domain (l+m≤1) for a wide variety of parameters. A consistently negative trace precludes the existence of closed orbits (limit cycles), thereby providing analytical support (but not proof) that membrane localization enhancement (DF > 1) is required for oscillations within our QSS approximation.

We next analyzed this two-variable system to test if having reaction rate enhancement from membrane localization for just a single enzyme, the kinase, was sufficient. Although the phosphatase can still bind and catalyze its substrate, it cannot do so from the 2D membrane surface, reducing the strength of the negative feedback. Setting the parameter corresponding to the recruitment of phosphatase P to the adaptor complex LpA to zero (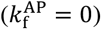) and consequently concentration LpAP to zero results in a trivial case in which L and Lp are in steady state and thus absent of oscillation across the domain. This analytically proves that negative feedback implemented via phosphatase recruitment by the adaptor is indispensable for generating oscillations in this QSS model structure. These results are summarized in Table 1, but we emphasize that while this analysis rigorously proves oscillations cannot occur under these specific limiting conditions (DF = 1 or 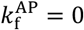) within the QSS domain of validity (SI), it does not constitute a proof of oscillation existence for the full model where these conditions are met. Proving the existence of limit cycles analytically for such complex nonlinear systems is often intractable. Therefore, we turn to extensive numerical simulations by using global optimization in parameter space. While numerical sampling cannot rule out oscillations, our sampling supports these analytical results that DF = 1 or 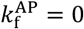 prevent oscillations (Table 1).

### III.4 Amplifying enzyme activation on the membrane dramatically expands the parameter space supporting oscillations

Based on our analysis of simplified models of our reaction network, the parameter *DF* = *V*/(*Ah*), which increases the rates that both enzymes bind their substrate, is a key factor in producing oscillations. A larger DF physically means that (i) the enzyme and substrate are concentrated in their now 2D search space (*V*/*A*) and/or (ii) the *h* lengthscale is small, indicating efficient 2D binding (but not diffusion-limited), collectively increasing the reactive flux to substrate binding. We performed stochastic optimization with a genetic algorithm to select for oscillatory individuals (or parameter sets) within a 17-dimensional space (DF+12 rates+4 initial concentrations, see Methods). When we fixed the DF to low values of 1-5, we found zero solutions, despite sampling millions of possible individuals with selection pressure for oscillations. When we increased DF to 6-10, we found 0.06% of individuals were oscillatory, but they occupied a highly restricted region of phenotype space characterized by low amplitudes and a narrow range of periods (Fig 2a). As DF increases into the 10-99 range and beyond, the accessible solution space expands dramatically, with the number of oscillating individuals increasing by orders-of-magnitude to represent ∼11% of individuals (Fig 2b). Phenotypically, we see oscillations with significantly higher amplitudes (approaching 1.0) become prevalent, and the range of achievable periods broadens considerably, spanning from tens to thousands of seconds, particularly at DF >100 (Fig 2b). This highlights DF, and by extension the physical parameter of cell or compartment geometry V/A, as a critical global modulator controlling the potential for oscillatory dynamics in this network architecture.

**Figure 2.**
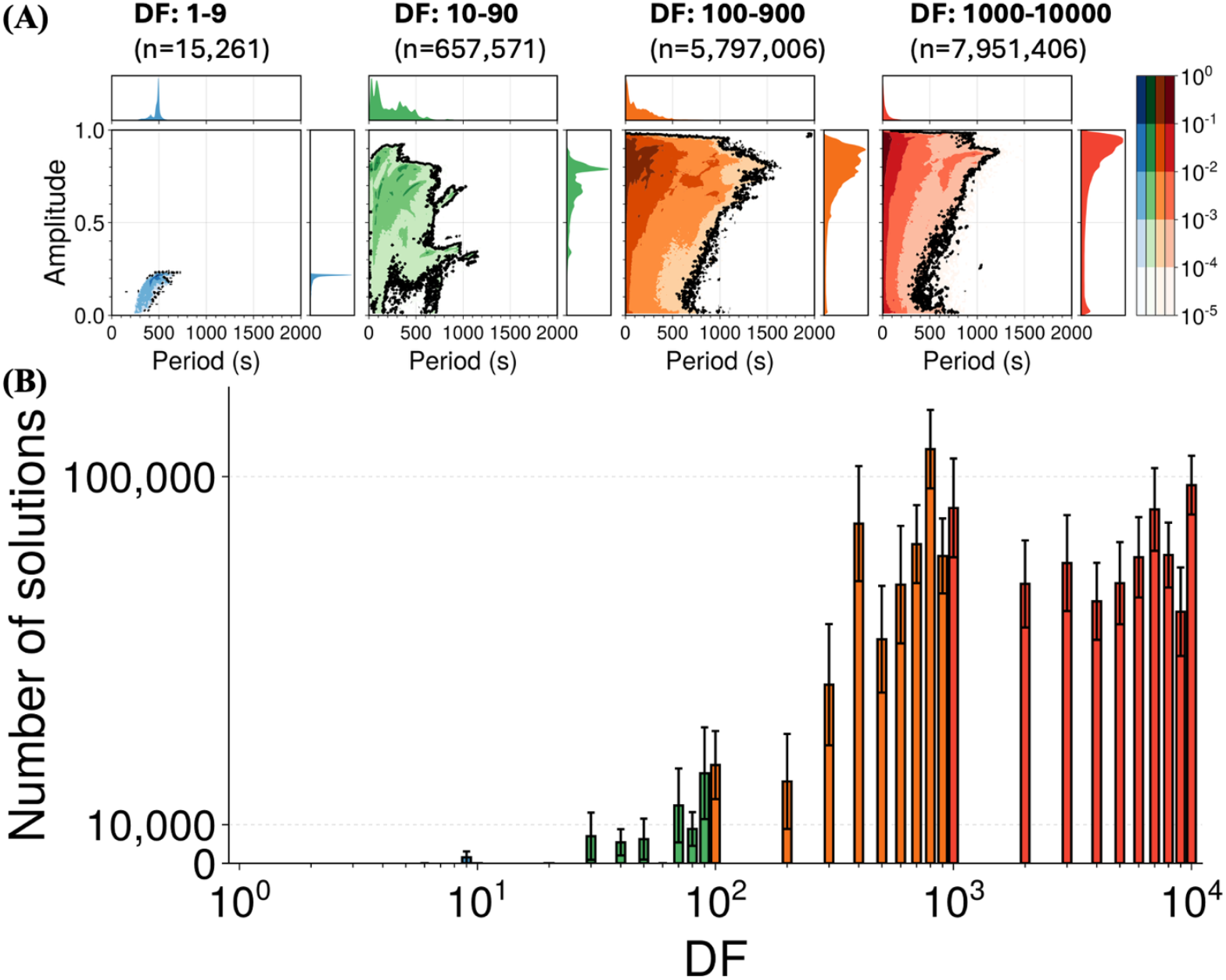
Amplifying enzyme activity via membrane localization with a higher DF Greatly Increases Both the Abundance and Phenotypic Diversity of Oscillatory Solutions. **(A)** Period-amplitude distributions for oscillatory parameter sets found across four distinct DF ranges (color-coded as in B). At low DF (blue, 1-9), viable solutions occupy a restricted phenotype space with low amplitudes and limited periods. As DF increases (green: 10-99, orange: 100-999, red: 1000-10000), the accessible region expands significantly, supporting oscillations with both wider period ranges and markedly higher amplitudes. Marginal histograms illustrate the increased diversity and range for both period (top) and amplitude (right) at higher DF values. **(B)** The number of distinct oscillatory individuals identified by genetic algorithm (mean ± SEM, n=10 independent runs) plotted against DF. For each DF value a single run can maximally produce 6×10^5^ individuals. Thus each DF range included sampling of 5.4 x 10^7^ parameter sets (9 bars x10 reps x6×10^5^ individuals/rep). A clear threshold effect is visible, with few solutions found below DF≈10. Above this threshold, the number of viable oscillatory solutions increases by over four orders of magnitude, saturating at high DF. This demonstrates that sufficient membrane localization drastically lowers the parametric requirements for achieving oscillations.

### III.5 Kinetic Constraints limit the Oscillatory Parameter Regime

While high DF values vastly expand the parameter space permissive for oscillations, we find that specific kinetic relationships remain crucial for establishing functional oscillatory circuits. To delineate these requirements, we analyzed the distributions of kinetic parameters within the identified oscillatory solution sets across different DF ranges (Fig 3). Our analysis reveals two key constraints governing the enzyme-substrate processing and adaptor-enzyme recruitment steps, both of which become less strict with higher DF. First, examining both enzyme-substrate interactions reveals a strong requirement for kinetic asymmetry. The Michaelis constant for the kinase 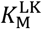 is always higher (or weaker) than the phosphatase 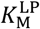 at the lowest DF values (Fig 3A). More explicitly, when we map the oscillatory solution space based on the ratio of catalytic rates 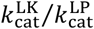 versus the ratio of forward binding rates 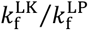. the low DF values require 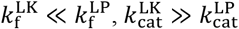 (Fig 3b). This corresponds to an *enzymatic efficiency asymmetry* where oscillations demand the kinase has slow substrate binding but fast catalysis, and the phosphatase has fast substrate binding but slow catalysis. The phosphatase must compete with the adaptor to bind Lp, and with the fast on-rate, it is then able to sequester Lp, preventing further recruitment of adaptor to the membrane. The slow catalysis then ensures that the kinase activity is also effectively slowed or blocked because its substrate is not being released. Once the substrate L is released, the kinase binds slowly, and this seems necessary to ensure the success of the negative feedback loop that returns A to 3D. While higher DF values significantly enlarge the permissible region in this parameter space, the bias towards the upper-left quadrant persists (Fig 3B), suggesting this asymmetry remains a beneficial, though less stringent, feature.

**Figure 3.**
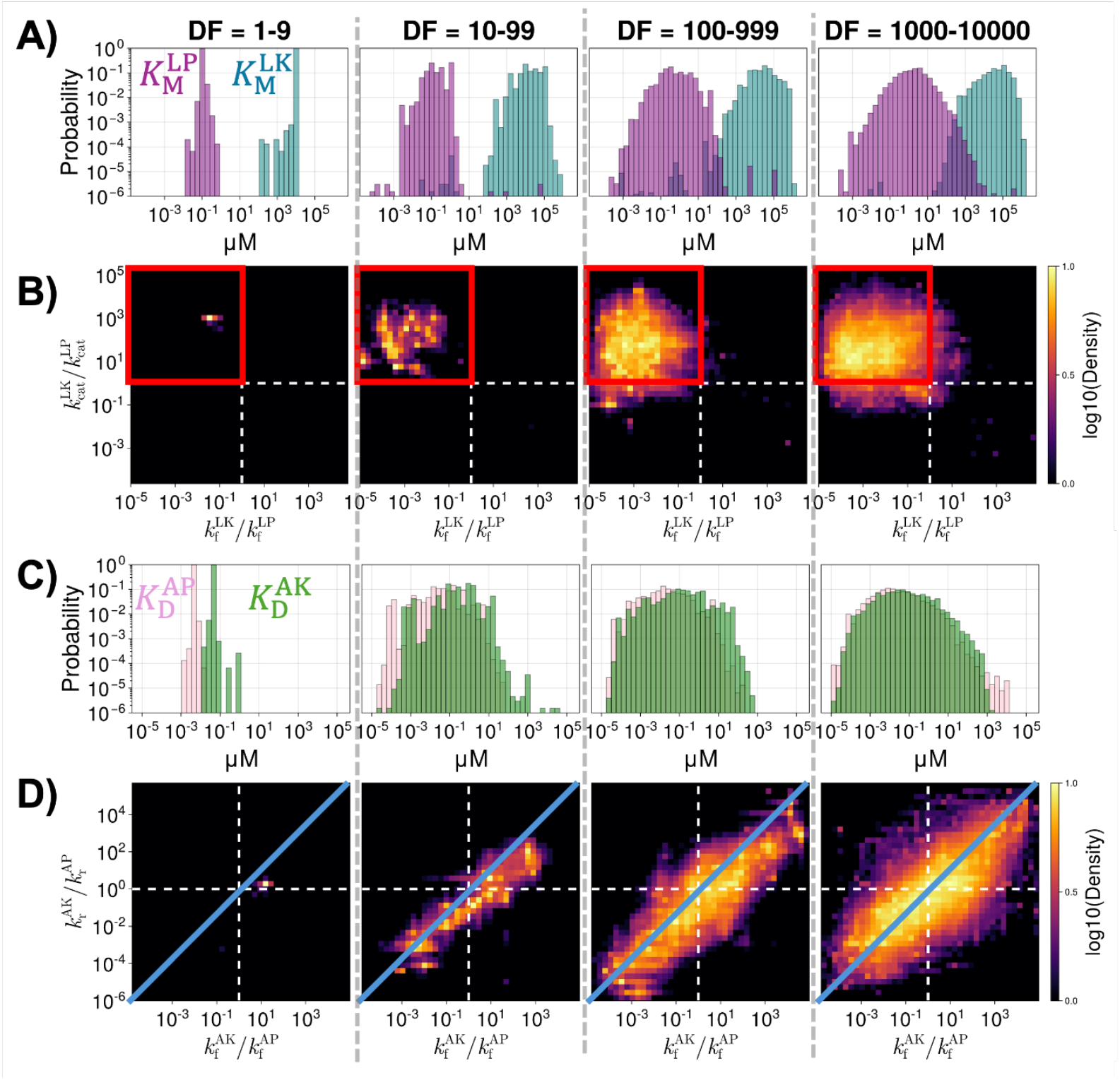
Oscillations rely on asymmetry in enzyme-substrate kinetics and relatively proportional adapter-enzyme binding. From left to right, each column shows parameters sampled at increasing DF ranges: DF=1-9; 10-100; 100-1000; and 1000-1000. (A) Constraints on enzyme-substrate processing kinetics. Distributions of Michaelis constants for the kinase (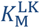, teal) are typically higher (weaker) than the phosphatase (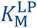, purple), and strictly so at the low DF values on the left column. B) 2D heatmaps showing the density of oscillatory solutions plotted by the ratio of enzyme-substrate catalytic rates 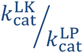 versus the ratio of enzyme-substrate forward binding rates 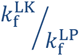. Solutions predominantly occupy the upper-left quadrant 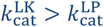 and 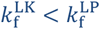 indicating a preference for *enzyme kinetic asymmetry*. The kinase is slow to bind and fast to catalyze its substrate, whereas the phosphatase is fast to bind and slow to catalyze its substrate. Higher DF values in right columns dramatically expand the permissible parameter region. (C) Constraints on adaptor-enzyme interactions: the distributions of dissociation constants for kinase-adaptor (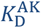. green) and phosphatase-adaptor (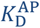, pink outline) binding show significant overlap, suggesting relatively balanced affinities and symmetry. D) 2D heatmaps showing solution density plotted by the ratio of adaptor-enzyme reverse rates 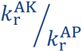 versus the ratio of adaptor-enzyme forward rates 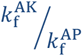. Solutions form a strong diagonal correlation, indicating *adaptor-enzyme binding proportionality*: the ratio of reverse rates is proportional to the ratio of forward rates 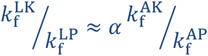.

**Figure 4.**
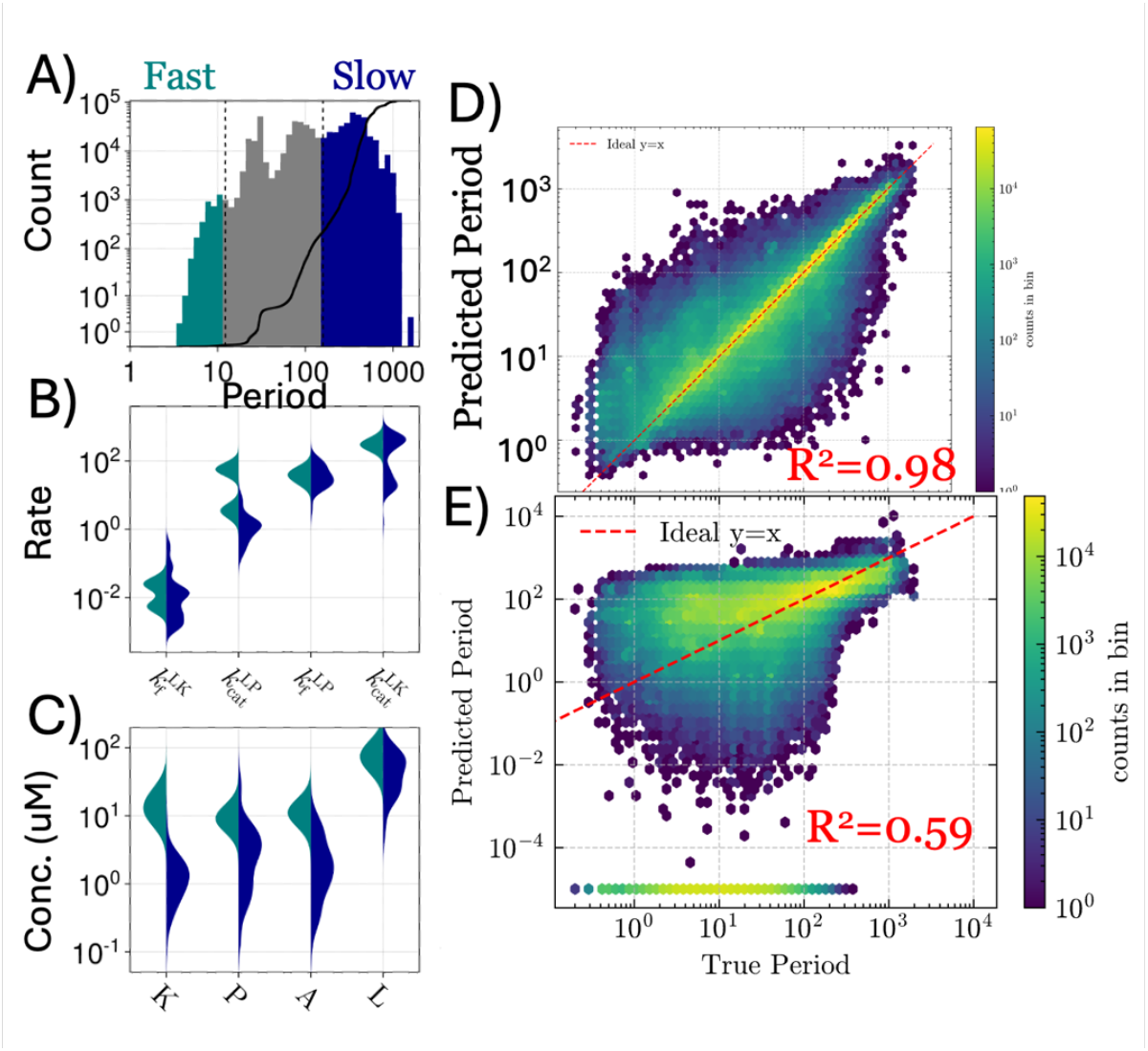
Period of oscillations correlate with phosphatase catalysis rate and enzyme-adaptor lifetimes. A) For our individuals in the DF range of 10-99, we split them into fast oscillators (short period-teal) and slow oscillators (large period-blue). B) The distributions of each parameter in these two sets of individuals shows the strongest divergence in the phosphatase catalysis, 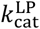, which is slow in slow oscillators. Enzyme forward rates and kinase catalysis are less strongly discriminating. C) Slow oscillators (blue) tend to have lower concentrations of total protein (K, A, P) and lipid (L) species. D) A multi-layer perceptron trained on our full dataset with an information bottleneck has very good accuracy in predicting continuous period values for each individual. E) Performing symbolic regression on the same dataset predicts the period with reasonable accuracy from a reduced subset of 6 input parameters, including 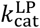.

Second, analyzing the kinetics of adaptor-enzyme recruitment reveals a distinct constraint based on proportionality (Fig 3C). The distributions of dissociation constants for kinase-adaptor 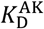 and phosphatase-adaptor 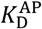 interactions exhibit considerable overlap across all DF ranges, suggesting that highly disparate affinities are disfavored (Fig 3C). More strictly, the heatmaps plotting the ratio of adaptor-enzyme dissociation rates 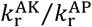 against the forward rates 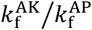 show solutions concentrated along a diagonal line (Fig 3D). This indicates a strong *adaptor-enzyme kinetics proportionality*, 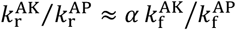, where *α* = 1, implying that the ratio of dissociation constants 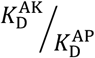 remains relatively constrained around 1 for viable solutions. The adaptor thus can have either strong or weak affinities to the enzymes, but it should maintain the same value for both enzymes. It is also common that both enzymes have widely varying unbinding rates 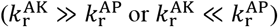, but this disparity in lifetimes must be compensated by equally disparate on-rates. Increasing DF broadens the diagonal band, allowing greater absolute variation in adaptor recruitment rates beyond strict proportionality.

### III.6 The period of oscillations trends with a subset of parameters

To go beyond establishing requirements for oscillations, we then interrogated what parameter regimes determine either fast (short period < 10 s) or slow (long period > 200s) oscillations. Despite that parameter combinations are inevitably coupled in this system (Fig 3), some individual parameters correlate fast vs slow periods when DF<100, with a persistent but less strong correlation at larger DF (Fig 4). The primary one is the catalysis rate of the phosphatase, 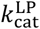, which is slower in slow oscillators (large periods). Further, slow oscillators have lower initial concentrations, particularly for the kinase K and adaptor A, even for larger DF values. Slow oscillators have stronger 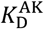 and 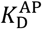 values for both adaptor-enzyme interactions. However, for the phosphatase, this stems from a faster on-rate, and for the kinase, from a slower off rate. Collectively, these trends indicate that slow periods arise in many cases from faster flux into the phosphatase state, and slower release of its substrate, along with a longer-lived interaction between the kinase and the adaptor, slowing transitions between states. Lower concentrations of all species will also slow reactive flux between states. The remaining parameters, including the rates of enzyme-substrate binding, showed limited correlation to period on their own.

To investigate period as a function of coupled parameters, we used machine-learning to demonstrate that a neural network can be trained to go beyond classification of the 17 input parameters and instead predict the specific period or oscillation with high accuracy. By creating a bottleneck in the NN, we improved its predictive performance and find that compression of the 17 variables to 2 latent variables has a similar predictive accuracy compared to retaining 17 latent variables. We used symbolic regression to try and predict the period from a single analytical function. While far from perfect, the symbolic function has reasonable predictive power, with R^2^=0.58, predicting the period as a function of 6 out of our 17 input parameters. For all variables normalized to reference values 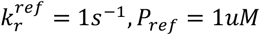, we find the normalized period as: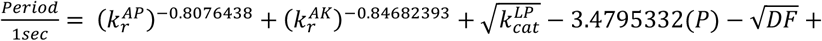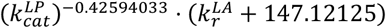. This function nicely recovers the trend from the single parameter analysis: a lower value of 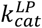 drives a slow oscillator/large period due to the highly weighted term: 147 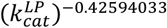. Similarly, fast dissociation and thus shorter lifetime of adaptor-enzyme interactions (large 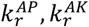), rather than enzyme-substrate interactions, drives faster oscillations (small period). Despite the obviously highly nonlinear relationship between our input parameters and the observed period of oscillation, our results suggest that a subset of key lifetime parameters tend to drive the speed of oscillations.

### III.7 Spatially-Resolved stochastic Simulations Validate Oscillatory Dynamics and Highlight Spatial Constraints

To assess the robustness of the oscillations to the assumptions of our deterministic ODE model, we compared its predictions against particle-based reaction-diffusion simulations with the NERDSS software ^32^. NERDSS simulations explicitly capture spatial organization, diffusion, and the inherent stochasticity arising from discrete molecular counts. The non-spatial ODE model captures the impact of membrane localization through the DF parameter, but assumes well-mixed conditions when in 3D, 2D and from 3D-2D. NERDSS incorporates coarse-grained molecular structure and orientation-dependent interactions into a particle-based reaction-diffusion framework. By including all microstates (intermediates) and only 1^st^ and 2^nd^ order reactions, our reaction networks are the same by design for NERDSS and the full ODE model, with NERDSS microscopic rates extracted from the macroscopic ODE rates and component diffusion coefficients (see Methods).

NERDSS simulations often successfully recapitulated the oscillatory behavior predicted by the deterministic ODE model for a variety of periods and amplitudes, albeit with increased noise as expected (Fig 5A). Despite the noise, NERDSS trajectories which successfully oscillate do so with a period and amplitude that are in general agreement with the smooth limit cycle produced by the ODEs (Fig 5b). Visual snapshots from the NERDSS simulation confirm the dynamic exchange of the adaptor A between the cytosol and the membrane, correlating with the peaks (high membrane localization) and troughs (peak delocalization) of the *A*_mem_ oscillation (Figure 6D insets). This concordance between the ODE and NERDSS results indicates that the core oscillatory mechanism derived from the ODE model, which incorporates dimensional reduction effects via DF, is robust and does not fundamentally break down when explicit spatial constraints and molecular noise are captured. This represents a key validation for future experimental tests, where space and stochasticity cannot be integrated out.

**Figure 5.**
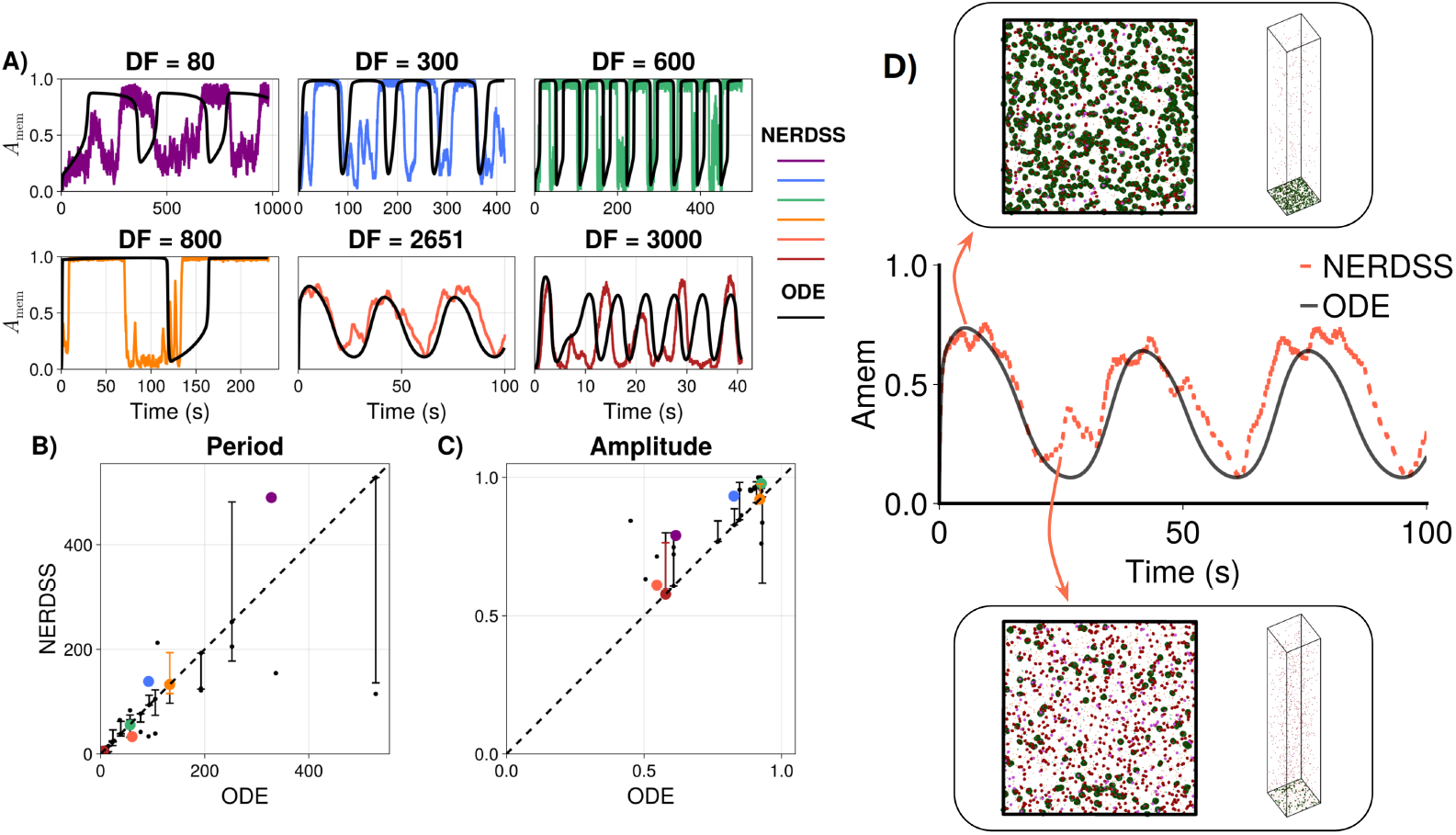
Spatially-resolved simulations validate the oscillatory dynamics predicted by the deterministic model. (A) Time courses of the membrane adaptor fraction (Amem) for a representative parameter set simulated at different Dimensionality Factor (DF) values. The trajectories from the spatially-resolved NERDSS model (colors, stochastic) qualitatively reproduce the oscillatory behavior predicted by the deterministic ODE model (black, smooth curves), despite inherent molecular noise. (B, C) Comparison of oscillation period (B) and amplitude (C) across a collection of oscillatory parameter sets. Each point plots the value derived from the ODE model against an approximate value measured from the corresponding NERDSS simulation. The data suggest a positive correlation for both metrics, with points distributed around the identity line (dashed). (D) Representative comparison of Amem dynamics from three simulation methodologies for a single parameter set: the deterministic ODE model (black line), the non-spatial Gillespie Stochastic Simulation Algorithm (SSA, dotted teal line), and the spatially-resolved NERDSS model (dashed orange line). The NERDSS trajectory, which explicitly models diffusion in 3D and 2D, shows closer agreement with the ODE prediction than the non-spatial SSA simulation does. Insets provide snapshots of the NERDSS simulation volume during peak adaptor localization (Top) and delocalization (Bottom), illustrating the physical basis of the oscillation.

**Figure 6.**
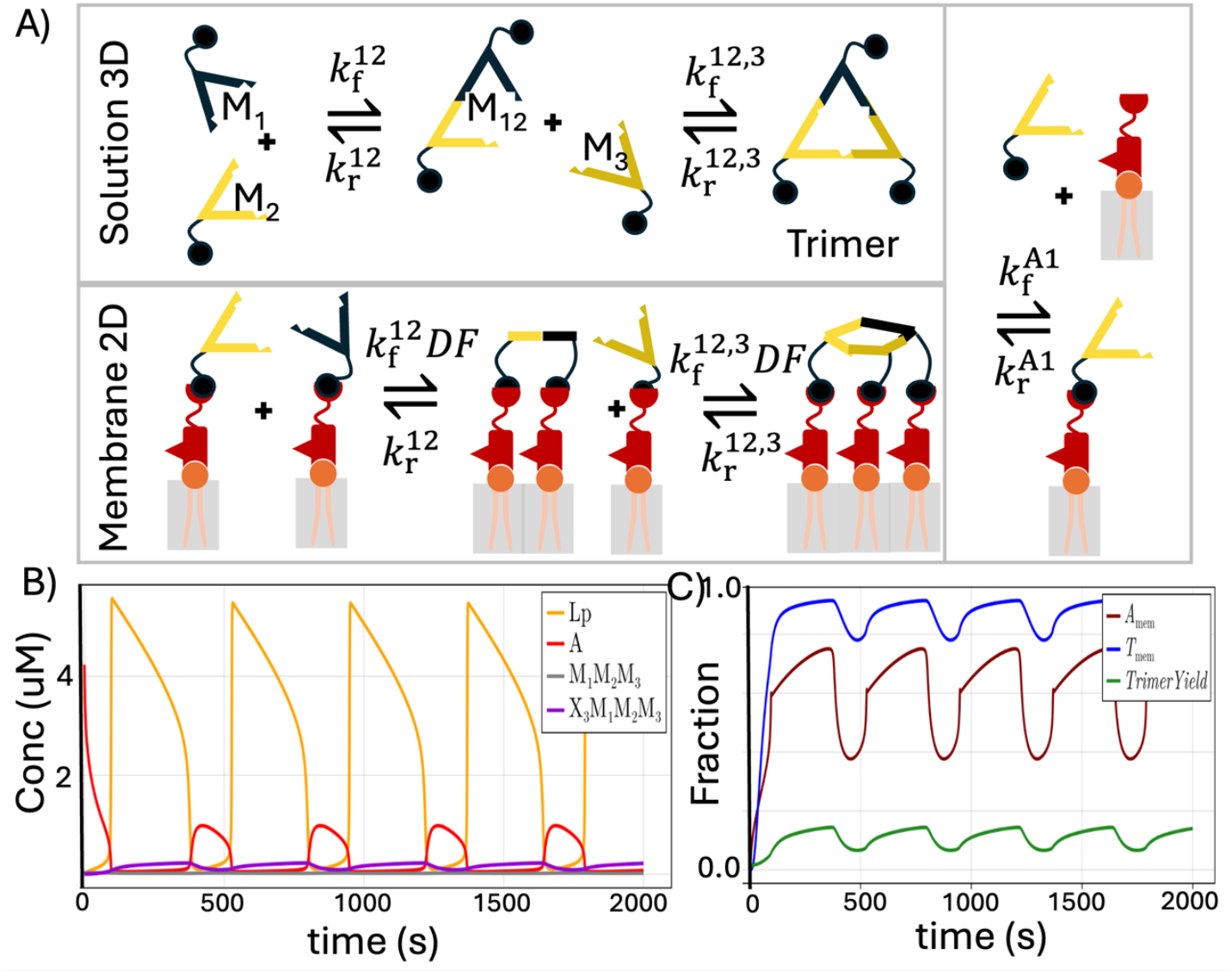
Membrane-Localized Oscillator drives oscillations in heterotrimer self-assembly. **(A)** Schematic illustrating the trimerizing subunits as they couple to the MemLo. Each distinct subunit of the heterotrimer has two interfaces for the trimerization, and a third interface to localize to the adaptor (right hand column). We illustrate here one possible assembly pathway in solution, although three are possible in purely 3D. The same pathways exist on the membrane, but scaled by the DF due to the 2D localization. Dimers and trimers can also localize to the membrane with only one or two adaptors bound. **(B)** One solution illustrates how the Lp species oscillates (yellow) driving oscillations in the solution (gray) and membrane-localized (purple) trimers. C) Our primary observable A_mem_ (brown) oscillates, creating simultaneous oscillations in the membrane-bound trimer normalized by total trimer (blue). The yield of trimer also oscillates as subunits move from the more dilute volume to the concentrated membrane (green).

Not all parameter sets we tested retained oscillations in the NERDSS simulations; approximately 72% of our randomly selected ODE-oscillatory parameter sets exhibited stable oscillations in NERDSS. This is despite the fact that we specifically tested individuals that were further from the diffusion limit for bimolecular association (*k*_f_ < 1*μM*^−1^*s*^−1^) to better accord with the well-mixed approximation^43^. This is particularly important for membrane localization reactions. Because the membrane exists only at one value of *z* in the simulation volume, diffusion to the surface can noticeably slow the effective on rate to the membrane for high *k*_f_ and/or high lipid copy numbers ^44^. We also initialized at least 100 copies of each monomer to minimize enhanced fluctuations that accompany small copy numbers. For systems that did not oscillate, increasing the copy numbers (via a larger V and A) did not fix the disagreement, indicating that the discrete abundances were not the primary issue. We speculate that the individuals where the deterministic oscillations relied on fast, DF-scaled bimolecular association rates exhibit highest discrepancies between the ODE predictions and NERDSS due to the limits of diffusional timescales.

### III.8 Self-assembly can be synchronized with oscillatory adaptor dynamics

A key potential application of synthetic biological oscillators is their use as programmable timing modules to control other cellular or biochemical processes. To explore the utility of the MemLO in this capacity, we established here that it could synchronize a canonical self-assembly reaction network, specifically the formation of a heterotrimer complex from three distinct monomer species, M_1_, M_2_, and M_3_ (Fig 6). Each heterotrimer subunit is endowed with an additional binding site for the adaptor A, and binding of A to these subunits does not compete with its pre-existing enzyme and lipid binding sites. This design creates parallel trimer assembly pathways: basal assembly occurring in the 3D cytosol, and enhanced assembly (for DF>1) when adaptor recruits subunits to the 2D membrane^26^. Critically, the assembly components do not need to be allosterically modified or post-translationally modified. Instead, the transition from weak assembly in 3D to stable assembly in 2D is driven by dimensional reduction ^45^, meaning that in principle, a wide variety of assembly subunits could be coupled to the MemLO core to drive cycles of assembly and disassembly.

In this case, the trimer is not purely moving on and off the membrane as the adaptor does. The monomeric subunits (and intermediates) move on and off the membrane and membrane localization stimulates assembly. Thus the yield of total trimer (relative to the maximum possible) also oscillates (Fig 6C). This simulation serves as a proof-of-concept that the MemLO’s periodicity can be exploited to drive oscillations in macromolecular self-assembly via adaptor-mediated membrane recruitment. It highlights the potential of using the MemLO, tuned by geometric factors (DF) as a dynamic scaffold to spatio-temporally orchestrate multi-protein complex formation.

## IV. DISCUSSION

In this study, we presented and analyzed a computational model for a novel class of membrane-localization oscillators (MemLOs), inspired by the lipid modification cycles involved in cellular processes like clathrin-mediated endocytosis. Our model demonstrates how the physical mechanism of membrane localization, coupled with specific network topology mediated by an adaptor protein A, can drive robust biochemical oscillations operating on rapid timescales (seconds to minutes), independent of slower transcription-translation processes. A key distinguishing feature of this MemLO design is the potential for oscillation frequency to be tuned directly by the system’s geometry, specifically the volume-to-surface area (V/A) ratio, offering a unique mode of control compared to traditional biochemical oscillators.

Our computational analyses revealed the fundamental principles underlying oscillation in this system. Our model enforces thermodynamic reversibility in all pairwise binding reactions, such that only the two enzyme catalysis reactions are irreversible, and mass is conserved. No explicit allostery is included. Instead, localization to the 2D membrane amplifies association reactions between enzymes and their substrates. We therefore established that the adaptor protein A is necessary to drive oscillations via binding to both enzymes. A sufficient enhancement of reactions on the 2D membrane (DF>1) also appears to be required for oscillations. High DF values, representing strong membrane localization effects, provide the necessary amplification or gain for the feedback loops, dramatically expanding the parameter space in which oscillations can occur and increasing the diversity of achievable oscillation phenotypes (periods and amplitudes).

Furthermore, we delineated a hierarchy of kinetic constraints defining the oscillatory regime. Successful oscillations rely not just on sufficient feedback gain but on a specific balance between kinetic parameters. This includes a preferred *enzymatic efficiency asymmetry* (contrasting kinetics for kinase vs. phosphatase substrate binding and catalysis) and an *adaptor-enzyme kinetics proportionality* (coupling the relative forward and reverse rates for adaptor-enzyme binding). The overall strictness of these kinetic requirements is substantially relaxed by high DF. A significant finding is the direct tunability of the oscillation period by DF, and thus by the physical V/A ratio. This presents a novel strategy for controlling biological timing without direct biochemical intervention. While we focused on DF, preliminary investigations suggest initial reactant concentrations may offer alternative tuning knobs, warranting further systematic study of the interplay between different control parameters and potential trade-offs between tunability and robustness.

The validity of the core mechanism in a more realistic setting was confirmed using spatially-resolved reaction-diffusion simulations (NERDSS). These simulations, explicitly accounting for space, diffusion, and molecular noise, successfully recapitulated the oscillations predicted by the non-spatial ODE model under a subset of parameter conditions. Finally, we demonstrated the potential utility of the MLO as a functional module by coupling it to a downstream heterotrimer self-assembly process via the adaptor protein A. The oscillator successfully synchronized the assembly pathway, driving periodic formation of the final trimer complex. This proof-of-concept illustrates how MLOs could be integrated into larger synthetic circuits to impose temporal control over diverse cellular processes, such as protein complex formation, signaling cascades, or timed release mechanisms.

Despite these promising computational and analytical findings, experimental validation remains the crucial next step. Constructing this MemLO system in vitro using purified components and synthetic vesicles, or potentially in vivo by repurposing existing cellular machinery (like components of the PIP cycle and associated adaptors), would be necessary to confirm the predicted oscillatory behavior and geometric tunability. Outstanding theoretical questions also remain, including a more comprehensive mapping of the parameter space governing tunability versus robustness, detailed analysis of noise effects in different regimes, and exploration of minimal network topologies capable of supporting membrane-localized oscillations. Overall, we think the memLo offers a promising avenues for synthetic biology applications ranging from dynamic biosensors to programmable timing elements in engineered cellular systems.

## ACKNOWLEDGMENTS

MEJ gratefully acknowledges funding from NIH MIRA award R35GM133644.

