## Supplemental Information for "A membrane-driven biochemical oscillator tunable by the volume to surface area ratio"

For

**Supplemental Methods and Text**

**Supplemental Figures**

**Supplemental Tables**

### SUPPLEMENTAL TEXT and METHODS

#### SI. Corresponding ODE model to capture dimensional reduction

To solve for this model nonspatially, such that all species can be converted between one another, we must choose what units we will use (or copy numbers) for concentrations. The equilibrium constant is in general a ratio of the forward and reverse rates: K_{eq}^{PS} =\ frac{k_{on}^{PS}}{k_{off}^{PS}}, where we leave out the dimension label for simplicity for all 3D reactions. The binding equilibrium for a species in 3D concentration, for example, \left[PS\right] = \frac{N_{PS}}{V} where N_{PS} are copy numbers of PS, is given by:

$$\frac{N_{PS}V}{N_P N_S} = \frac{k_{on}^{PS}}{k_{\text{off}}^{PS}} \tag{1}$$

In these 3D units, the association rate has units of \frac{V}Error! Bookmark not defined. and dissociation has units of \frac{1}Error! Bookmark not defined.. For the 2D reactions, however, we have that

$$\frac{N_{PSN}A}{N_{PN}N_S} = \frac{k_{on}^{2D,PS}}{k_{\text{off}}^{2D,PS}} \tag{2}$$

Where association and dissociation rates have units of $\frac{\mathrm{V}}{\mathrm{time}}$ and $\frac{1}{\mathrm{time}}$ respectively. It is important to note that it is not true that $\frac{\mathrm{N_{PSN}V}}{\mathrm{N_{PN}N_S}} = \frac{\mathrm{k_{on}^{3D,PS}}}{\mathrm{k_{off}^{3D,PS}}}$, as this relation asserts that the copies are performing a 3D search in a volume $\mathrm{V}$. Instead, to put the Eq. 4 in volume units, they must be re-scaled by multiplying both sides by the dimensionality factor:

$$DF = \frac{V}{Ah} \tag{3}$$

The rates are related to one another via

$$k_{on}^{2D} = \frac{k_{on}^{3D}}{h'} \tag{4a}$$

$$k_b^{2D} = ck_b^{3D} \tag{4b}$$

where {h^\prime} has units of area, and the off-rates can be distinct in 3D vs 2D by a scalar, $c$. The equilibrium constants are thus related by $h = ch'$,

$$K_{eq}^{2D} = \frac{K_{eq}^{3D}}{h} \tag{5}$$

By multiplying both sides in Eq. 4 by $DF$ and using Eq. 6 and 7, we find:

$$\frac{N_{PSN}V}{N_{PN}N_S} = \frac{Vk_{on}^{3D,PS}}{Ahk_{\text{off}}^{3D,PS}} = DF \cdot K_{eq}^{3D,PS} \tag{6}$$

In this way, we can see that the bimolecular equilibria between the 2D species is related to the 3D equilibrium constant and the 3D concentrations via the factor $DF$.

Equivalently, we know that for the bimolecular association reaction in 1D,

$$\frac{dN_{PSN}}{Ldt} = -\frac{N_{PSN}k_{off}^{2D,PS}}{V} + \frac{k_{on}^{2D,PS}N_{PN}N_S}{V^2} \tag{7}$$

If we multiply both sides of Eq. 7 by \frac{Ah}{V}, we get:

$$\frac{dN_{PSN}}{Vdt} = -\frac{N_{PSN}ck_{off}^{3D,PS}}{V} + \frac{ck_{on}^{3D,PS}N_{PN}N_S}{VAh} \tag{8}$$

Finally, this equation will be solved for in fully 3D units by multiplying the 3rd term by \frac{V}{V}, giving

$$\frac{dN_{PSN}}{Vdt} = -\frac{N_{PSN}ck_{off}^{3D,PS}}{V} + \frac{DF \cdot ck_{on}^{3D,PS}N_{PN}N_S}{V^2} \tag{9}$$

Which is exactly the equation in volume units, with the association step multiplied by the $DF$, and we allow a rescaling between 3D and 2D of the dissociation rate as well, noting that the $DF$ used here is based on the equilibrium constant Eq. 7, not the forward rate, hence the factor of $c$ also showing up in the association rate.

**SI.1 Reaction Network for full, unrestricted model.** There are 16 species in the full model, or unrestricted model, interacting via 25 reactions. All interfaces can bind in all intermediate states, and the model is fully compatible with NERDSS. The 5 monomeric species are A,K,P,L and Lp. A, K, and P each have two distinct binding sites, one to bind a protein (for A, both K and P bind at the same position), and one to bind a lipid. L and Lp each have a single interface for binding a protein. This gives rise to 5 dimer species: AK, AP, LpA, LpP, LK, that form via 5 reversible 3D reactions. The dimers can bind to monomers to produce 4 distinct 3-mers: LpAK,

LpAP, AKL, and APLp via 8 reversible 3D reactions. Finally, dimer-dimer reactions (2rxns), or trimer-monomer reactions (4 rxns) can produce the two 4-mer species: LpAKL and LpAPLp. There are two distinct catalysis reactions, one for K and one for P. They are performed by the species LK, AKL, LpAKL, and LpP, LpPA, LpPALp, for 6 additional reactions, totaling 25. The pairwise reactions are listed below with their reaction rates. All 2D reactions are listed in Volume units by rescaling via the dimensionality factor.

$$\mathrm{Lp} + \mathrm{A} \underset{k_r^{\mathrm{LA}}}{\overset{k_f^{\mathrm{LA}}}{\rightleftharpoons}} \mathrm{LpA} \tag{1}$$

$$\mathrm{L} + \mathrm{K} \underset{k_r^{\mathrm{LK}}}{\overset{k_f^{\mathrm{LK}}}{\rightleftharpoons}} \mathrm{LK} \xrightarrow{k_{cat}^{\mathrm{LK}}} \mathrm{Lp} + \mathrm{K} \tag{2}$$

$$\mathrm{Lp} + \mathrm{P} \underset{k_r^{\mathrm{LP}}}{\overset{k_f^{\mathrm{LP}}}{\rightleftharpoons}} \mathrm{LpP} \xrightarrow{k_{cat}^{\mathrm{P}}} \mathrm{L} + \mathrm{P} \tag{3}$$

$$\mathrm{LpA} + \mathrm{K} \underset{k_r^{\mathrm{AK}}}{\overset{k_f^{\mathrm{AK}}}{\rightleftharpoons}} \mathrm{LpAK} \tag{4}$$

$$\mathrm{LpAK} + \mathrm{L} \underset{k_r^{\mathrm{LK}}}{\overset{DF \cdot k_f^{\mathrm{LK}}}{\rightleftharpoons}} \mathrm{LpAKL} \xrightarrow{k_{cat}^{\mathrm{L}}} \mathrm{Lp} + \mathrm{LpAK} \tag{5}$$

$$\mathrm{LpA} + \mathrm{P} \underset{k_r^{\mathrm{AP}}}{\overset{k_f^{\mathrm{AP}}}{\rightleftharpoons}} \mathrm{LpAP} \tag{6}$$

$$\mathrm{Lp} + \mathrm{LpAP} \underset{k_r^{\mathrm{LP}}}{\overset{DF \cdot k_f^{\mathrm{LP}}}{\rightleftharpoons}} \mathrm{LpAPLp} \xrightarrow{k_{cat}^{\mathrm{Lp}}} \mathrm{L} + \mathrm{LpAP} \tag{7}$$

$$\mathrm{Lp} + \mathrm{AK} \underset{k_r^{\mathrm{LA}}}{\overset{k_f^{\mathrm{LA}}}{\rightleftharpoons}} \mathrm{LpAK} \tag{8}$$

$$\mathrm{Lp} + \mathrm{AKL} \underset{k_r^{\mathrm{LA}}}{\overset{DF \cdot k_f^{\mathrm{LA}}}{\rightleftharpoons}} \mathrm{LpAKL} \tag{9}$$

$$\mathrm{Lp} + \mathrm{AP} \underset{k_r^{\mathrm{LA}}}{\overset{k_f^{\mathrm{LA}}}{\rightleftharpoons}} \mathrm{LpAP} \tag{10}$$

$$\mathrm{Lp} + \mathrm{APLp} \underset{k_r^{\mathrm{LA}}}{\overset{DF \cdot k_f^{\mathrm{LA}}}{\rightleftharpoons}} \mathrm{LpAPLp} \tag{11}$$

$$\mathrm{A} + \mathrm{K} \underset{k_r^{\mathrm{AK}}}{\overset{k_f^{\mathrm{AK}}}{\rightleftharpoons}} \mathrm{AK} \tag{12}$$

$$\mathrm{A} + \mathrm{P} \underset{k_r^{\mathrm{AP}}}{\overset{k_f^{\mathrm{AP}}}{\rightleftharpoons}} \mathrm{AP} \tag{13}$$

$$\mathrm{A} + \mathrm{LK} \underset{k_r^{\mathrm{AK}}}{\overset{k_f^{\mathrm{AK}}}{\rightleftharpoons}} \mathrm{AKL} \tag{14}$$

$$\mathrm{A} + \mathrm{LpP} \underset{k_r^{\mathrm{AP}}}{\overset{k_f^{\mathrm{AP}}}{\rightleftharpoons}} \mathrm{APLp} \tag{15}$$

$$\mathrm{LpA} + \mathrm{LK} \underset{k_r^{\mathrm{AK}}}{\overset{DF \cdot k_f^{\mathrm{AK}}}{\rightleftharpoons}} \mathrm{LpAKL} \tag{16}$$

$$\mathrm{LpA} + \mathrm{LpP} \underset{k_r^{\mathrm{AP}}}{\overset{DF \cdot k_f^{\mathrm{AP}}}{\rightleftharpoons}} \mathrm{LpAPLp} \tag{17}$$

$$\mathrm{AK} + \mathrm{L} \underset{k_r^{\mathrm{LK}}}{\overset{k_f^{\mathrm{LK}}}{\rightleftharpoons}} \mathrm{AKL} \xrightarrow{k_{cat}^{\mathrm{LK}}} \mathrm{Lp} + \mathrm{AK} \tag{18}$$

$$\mathrm{AP} + \mathrm{Lp} \underset{k_r^{\mathrm{LP}}}{\overset{k_f^{\mathrm{LP}}}{\rightleftharpoons}} \mathrm{APLp} \xrightarrow{k_{cat}^{\mathrm{P}}} \mathrm{L} + \mathrm{AP} \tag{19}$$

### SI.2 Ordinary Differential Equations for the full 16 variable model:

$$\frac{\mathrm{d}L(t)}{\mathrm{d}t} = kcat^{LP}\mathrm{LpP}(t) + kcat^{LP}\mathrm{APLp}(t) + kcat^{LP}\mathrm{LpAPLp}(t) + kr^{LK}\mathrm{AKL}(t) + kr^{LK}\mathrm{LpAKL}(t) + kr^{LK}\mathrm{LK}(t) - kf^{LK}K(t)L(t) - kf^{LK}\mathrm{AK}(t)L(t) - DFkf^{LK}\mathrm{LpAK}(t)L(t)$$

$$\frac{\mathrm{d}K(t)}{\mathrm{d}t} = kcat^{LK}\mathrm{LK}(t) + kr^{AK}\mathrm{LpAK}(t) + kr^{AK}\mathrm{AK}(t) + kr^{LK}\mathrm{LK}(t) - kf^{AK}\mathrm{LpA}(t)K(t) - kf^{AK}A(t)K(t) - kf^{LK}K(t)L(t)$$

$$\frac{\mathrm{d}P(t)}{\mathrm{d}t} = kcat^{LP}\mathrm{LpP}(t) + kr^{AP}\mathrm{AP}(t) + kr^{AP}\mathrm{LpAP}(t) + kr^{LP}\mathrm{LpP}(t) - kf^{AP}\mathrm{LpA}(t)P(t) - kf^{AP}A(t)P(t) - kf^{LP}\mathrm{Lp}(t)P(t)$$

$$\frac{\mathrm{d}A(t)}{\mathrm{d}t} = kr^{AK}\mathrm{AKL}(t) + kr^{AK}\mathrm{AK}(t) + kr^{AP}\mathrm{AP}(t) + kr^{AP}\mathrm{APLp}(t) + kr^{LA}\mathrm{LpA}(t) - kf^{AK}A(t)\mathrm{LK}(t) - kf^{AK}A(t)K(t) - kf^{AP}\mathrm{LpP}(t)A(t) - kf^{AP}A(t)P(t) - kf^{LA}\mathrm{Lp}(t)A(t)$$

$$\frac{\mathrm{dLp}(t)}{\mathrm{d}t} = kcat^{LK}\mathrm{AKL}(t) + kcat^{LK}\mathrm{LpAKL}(t) + kcat^{LK}\mathrm{LK}(t) + kr^{LA}\mathrm{LpAP}(t) + kr^{LA}\mathrm{LpAPLp}(t) + kr^{LA}\mathrm{LpAKL}(t) + kr^{LA}\mathrm{LpAK}(t) + kr^{LA}\mathrm{LpA}(t) + kr^{LP}\mathrm{LpP}(t) + kr^{LP}\mathrm{APLp}(t) + kr^{LP}\mathrm{LpAPLp}(t) - kf^{LA}\mathrm{AP}(t)\mathrm{Lp}(t) - kf^{LA}\mathrm{Lp}(t)A(t) - kf^{LA}\mathrm{Lp}(t)\mathrm{AK}(t) - kf^{LP}\mathrm{AP}(t)\mathrm{Lp}(t) - kf^{LP}\mathrm{Lp}(t)P(t) - DFkf^{LA}\mathrm{APLp}(t)\mathrm{Lp}(t) - DFkf^{LA}\mathrm{AKL}(t)\mathrm{Lp}(t) - DFkf^{LP}\mathrm{LpAP}(t)\mathrm{Lp}(t)$$

$$\frac{\mathrm{dLpA}(t)}{\mathrm{d}t} = kr^{AK}\mathrm{LpAKL}(t) + kr^{AK}\mathrm{LpAK}(t) + kr^{AP}\mathrm{LpAP}(t) + kr^{AP}\mathrm{LpAPLp}(t) - kr^{LA}\mathrm{LpA}(t) - kf^{AK}\mathrm{LpA}(t)K(t) - kf^{AP}\mathrm{LpA}(t)P(t) + kf^{LA}\mathrm{Lp}(t)A(t) - DFkf^{AK}\mathrm{LpA}(t)\mathrm{LK}(t) - DFkf^{AP}\mathrm{LpP}(t)\mathrm{LpA}(t)$$

$$\frac{\mathrm{dLK}(t)}{\mathrm{d}t} = -kcat^{LK}\mathrm{LK}(t) + kr^{AK}\mathrm{AKL}(t) + kr^{AK}\mathrm{LpAKL}(t) - kr^{LK}\mathrm{LK}(t) - kf^{AK}A(t)\mathrm{LK}(t) + kf^{LK}K(t)L(t) - DFkf^{AK}\mathrm{LpA}(t)\mathrm{LK}(t)$$

$$\frac{\mathrm{dLpP}(t)}{\mathrm{d}t} = -kcat^{LP}\mathrm{LpP}(t) + kr^{AP}\mathrm{APLp}(t) + kr^{AP}\mathrm{LpAPLp}(t) - kr^{LP}\mathrm{LpP}(t) - kf^{AP}\mathrm{LpP}(t)A(t) + kf^{LP}\mathrm{Lp}(t)P(t) - DFkf^{AP}\mathrm{LpP}(t)\mathrm{LpA}(t)$$

$$\frac{\mathrm{dLpAK}(t)}{\mathrm{d}t} = kcat^{LK}\mathrm{LpAKL}(t) - kr^{AK}\mathrm{LpAK}(t) - kr^{LA}\mathrm{LpAK}(t) + kr^{LK}\mathrm{LpAKL}(t) + kf^{AK}\mathrm{LpA}(t)K(t) + kf^{LA}\mathrm{Lp}(t)\mathrm{AK}(t) - DFkf^{LK}\mathrm{LpAK}(t)L(t)$$

$$\frac{\mathrm{dLpAP}\,(t)}{\mathrm{d}t} = kcat^{LP}\mathrm{LpAPLp}\,(t) - kr^{AP}\mathrm{LpAP}\,(t) - kr^{LA}\mathrm{LpAP}\,(t) + kr^{LP}\mathrm{LpAPLp}\,(t) + kf^{AP}\mathrm{LpA}\,(t)\,P\,(t) + kf^{LA}\mathrm{AP}\,(t)\,\mathrm{Lp}\,(t) - DFkf^{LP}\mathrm{LpAP}\,(t)\,\mathrm{Lp}\,(t)$$

$$\frac{\mathrm{dLpAKL}\,(t)}{\mathrm{d}t} = -kcat^{LK}\mathrm{LpAKL}\,(t) - kr^{AK}\mathrm{LpAKL}\,(t) - kr^{LA}\mathrm{LpAKL}\,(t) - kr^{LK}\mathrm{LpAKL}\,(t) + DFkf^{AK}\mathrm{LpA}\,(t)\,\mathrm{LK}\,(t) + DFkf^{LA}\mathrm{AKL}\,(t)\,\mathrm{Lp}\,(t) + DFkf^{LK}\mathrm{LpAK}\,(t)\,L\,(t)$$

$$\frac{\mathrm{dLpAPLp}\,(t)}{\mathrm{d}t} = -kcat^{LP}\mathrm{LpAPLp}\,(t) - kr^{AP}\mathrm{LpAPLp}\,(t) - kr^{LA}\mathrm{LpAPLp}\,(t) - kr^{LP}\mathrm{LpAPLp}\,(t) + DFkf^{AP}\mathrm{LpP}\,(t)\,\mathrm{LpA}\,(t) + DFkf^{LA}\mathrm{APLp}\,(t)\,\mathrm{Lp}\,(t) + DFkf^{LP}\mathrm{LpAP}\,(t)\,\mathrm{Lp}\,(t)$$

$$\frac{\mathrm{dAK}\,(t)}{\mathrm{d}t} = kcat^{LK}\mathrm{AKL}\,(t) - kr^{AK}\mathrm{AK}\,(t) + kr^{LA}\mathrm{LpAK}\,(t) + kr^{LK}\mathrm{AKL}\,(t) + kf^{AK}A\,(t)\,K\,(t) - kf^{LA}\mathrm{Lp}\,(t)\,\mathrm{AK}\,(t) - kf^{LK}\mathrm{AK}\,(t)\,L\,(t)$$

$$\frac{\mathrm{dAP}\,(t)}{\mathrm{d}t} = kcat^{LP}\mathrm{APLp}\,(t) - kr^{AP}\mathrm{AP}\,(t) + kr^{LA}\mathrm{LpAP}\,(t) + kr^{LP}\mathrm{APLp}\,(t) + kf^{AP}A\,(t)\,P\,(t) - kf^{LA}\mathrm{AP}\,(t)\,\mathrm{Lp}\,(t) - kf^{LP}\mathrm{AP}\,(t)\,\mathrm{Lp}\,(t)$$

$$\frac{\mathrm{dAKL}\,(t)}{\mathrm{d}t} = -kcat^{LK}\mathrm{AKL}\,(t) - kr^{AK}\mathrm{AKL}\,(t) + kr^{LA}\mathrm{LpAKL}\,(t) - kr^{LK}\mathrm{AKL}\,(t) + kf^{AK}A\,(t)\,\mathrm{LK}\,(t) + kf^{LK}\mathrm{AK}\,(t)\,L\,(t) - DFkf^{LA}\mathrm{AKL}\,(t)\,\mathrm{Lp}\,(t)$$

$$\frac{\mathrm{dAPLp}\,(t)}{\mathrm{d}t} = -kcat^{LP}\mathrm{APLp}\,(t) - kr^{AP}\mathrm{APLp}\,(t) + kr^{LA}\mathrm{LpAPLp}\,(t) - kr^{LP}\mathrm{APLp}\,(t) + kf^{AP}\mathrm{LpP}\,(t)\,A\,(t) + kf^{LP}\mathrm{AP}\,(t)\,\mathrm{Lp}\,(t) - DFkf^{LA}\mathrm{APLp}\,(t)\,\mathrm{Lp}\,(t)$$

### SI.3 Mass conservation in the unrestricted/full model

Given the 16 species defined above, the mass conservation can be written as:

Our system of ODEs has 16 time-dependent variables interacting via 19 reversible reactions and 6 catalytic reactions. Monomers: A, K, P, L, Lp, dimers: AK, AP, LpA, LpP, LK, three-mers: LpAK, LpAP, AKL, and APLp, and 4-mers LpAPLp and LpAKL represent the concentrations of the complexed and noncomplexed forms of the chemical species in our model. Four mass conservation laws constrain the fixed total concentrations of lipid, adaptor protein, kinase, and phosphatase.

$$Lipid_{total} = L + Lp + LK + LpP + LpA + LpAK + LpAP + 2LpAKL + 2LpAPLp + APLp + AKL \quad (4a)$$

$$A_{total} = A + LpA + LpAK + LpAP + LpAKL + LpAPLp + AK + AP + APLp + AKL \quad (4b)$$

$$K_{total} = K + LK + LpAK + LpAKL + AK + AKL \quad (4c)$$

$$P_{total} = P + LpP + LpAP + LpAPLp + AP + APLp \quad (4d)$$

**SI.4 Thermodynamic Cycles in the full model:**

For several of our higher-order (3-mer and 4-mer species), there is more than one binding pathway that can reach them. Although we have irreversible catalytic reactions in our network, the pathways that are reversible must maintain energy conservation, and this constrains the free energies and thus the relative reaction rates for these processes. Our models are defined to be energy conserving around these cycles, by choosing the same value of *h* for all 2D reactions, and using the same reaction rates when the same interfaces are involved. While this is not the only choice, it is the simplest, with the fewest parameters. It means that the products of the $K_D$s as we move around the cycle is identical in both directions, and thus no energy is created or destroyed by returning around the cycle to the start point.

Example: A<->LpA (+Lp) ; LpA<->LpAP (+P)

A<->AP (+P) ; AP<->LpAP (+Lp).

Example 2 (with DF): LpA <->LpAP (+P); LpAP <->LpAPLp (DF+Lp)

LpA<->LpAPLp (DF + PLp).

**SII.1 Reaction Network for the simplified, restricted model.** There are 12 species in the restricted model, interacting via 11 reactions. This model has been simplified to remove intermediates that are not key driving forces for producing oscillations. We have the same 5 monomeric species as the full model, A,K,P,L and Lp, and they have the same number of interfaces, but we restrict some of the binding reactions involving the adaptor A, eliminating 4 species. Specifically, we assume that A is inhibited from binding enzymes in solution (from 3D), and therefore can only bind either P or K after it binds to the lipids. We thus produce 3 dimer species: LpA, LpP, LK, that form via 3 reversible 3D reactions. The dimers can bind to monomers to produce 2 distinct 3-mers: LpAK, LpAP, via 2 reversible 3D reactions. Here we have enforced that A cannot bind to the LK or LpP species, because it is still in solution. Finally, trimer-monomer reactions (2 rxns) can produce the two 4-mer species: LpAKL and LpAPLp. There are two distinct catalysis reactions, one for K and one for P. They are performed by the species LK, LpAKL, and LpP, LpPALp, for 4 additional reactions, totaling 11. The pairwise reactions are listed below with their reaction rates. All 2D reactions are listed in Volume units by rescaling via the dimensionality factor. We still have the same number of 12 reaction rate constants, as dimerization reactions are still occurring between adaptor and enzymes, but requiring the LpA version of the adaptor.

**SII.2 The ODEs are given by:**

\\mathbf{mathbfmathbfmathbfdL}/\\mathbf{mathbfmathbfmathbfdt} =\mathbit{kca}\mathbit{t}^{\mathbit{L}\ \mathbit{P}}

**SII.3 Mass conservation in the restricted model.**

$$\mathrm{Lipid_{total}} = \mathrm{L} + \mathrm{Lp} + \mathrm{LK} + \mathrm{LpP} + \mathrm{LpA} + \mathrm{LpAK} + \mathrm{LpAP} + 2\mathrm{LpAKL} + 2\mathrm{LpAPLp} \tag{4a}$$

$$\mathrm{A_{total}} = \mathrm{A} + \mathrm{LpA} + \mathrm{LpAK} + \mathrm{LpAP} + \mathrm{LpAKL} + \mathrm{LpAPLp} \tag{4b}$$

$$\mathrm{K_{total}} = \mathrm{K} + \mathrm{LK} + \mathrm{LpAK} + \mathrm{LpAKL} \tag{4c}$$

$$\mathrm{P_{total}} = \mathrm{P} + \mathrm{LpP} + \mathrm{LpAP} + \mathrm{LpAPLp} \tag{4d}$$

**PARAMETER REGIMES FOR CONCENTRATIONS: L>A :** Typically we will want the concentration of L to be higher than A. Otherwise, if A>>L, then it is impossible for a large fraction of A to transition to the membrane, because it must have enough binding sites. So we can argue that we are generally interested in regimes where we can get relatively large amplitude oscillations as measured by A going on and off the membrane, and therefore we will have L>A. In real systems, enzymes have much lower concentrations than A, and it would likely be more energy efficient if we could have large amplitude oscillations with small numbers of enzymes, so if A>K in particular is true, and yet we can get large amplitudes, that should be more efficient than the same size oscillations with K>A (because I'm assuming there would be more L->Lp conversions in the latter). K is the one that consumes ATP. Phosphatases use hydrolysis to break bonds.

**GENETIC ALGORITHM:**

For our parameter sweep, we had to choose certain hyperparameter values for our GA that would maximize both the quality of saved individuals and coverage of the search space. These hyperparameters were the population size, crossover rate, mutation rate, mutation scalar (explained in the mutation section), and number of tournament groups the population would be divided into for selection. We conducted a hyperparameter sweep via grid search where we varied a single hyperparameter while holding all others at nominal values, using the same seed to control for the stochasticity of the GA.

The results of the hyperparameter sweep are shown in Figure S2, where the performance metrics were max-pairwise distance in the final population to measure search coverage, and average fitness of the final population to measure search quality. Our conclusion from this analysis was that population size was by far the most important hyperparameter and dominated the effect of any variation in the other hyperparameters. Therefore, we chose a population size of 100,000, which was the maximum feasible size within our computational constraints.

The initial population P is generated by sampling 100000 genotype vectors $g_j = [k_{j,1}\ k_{j,2}\ \cdots\ k_{j,n}]$ where each gene element of genotype $g_j$ is sampled log-uniformly:

$$k_{j,i} = 10^{Uniform(log_{10}(min\,(k),\ log_{10}(max\,(k))} \quad \text{for } i = 1, 2, \ldots, n \text{ and } j = 1, 2, \ldots, m \tag{8a}$$

and concatenated into the population matrix M

$$\mathrm{M} = \begin{bmatrix} k_{1,1} & \cdots & k_{1,n} \\ \vdots & \ddots & \vdots \\ k_{m,1} & \cdots & k_{m,n} \end{bmatrix} \tag{8b}$$

where n is the number of genes in a genotype, $\mathrm{l}b_i$ and $\mathrm{ub}_i$ are the lower and upper bounds for i-th gene of a genotype, and $\mathrm{m} = 100000$ for the number of genotype vectors in the population.

M is then transformed with mutation, crossover, and selection operations for every generation, for 5 generations total (not including the initialization).

***Mutation***

The chromosomal mutation rate (whether a vector is chosen to be mutated) was set high at 95% in order to encourage exploration and maintain population diversity.

If mutation is performed, we use the Polynomial Mutation (PLM) scheme, which is mathematically represented as:

$$c_i' = c_i + \Delta \cdot \delta_i \tag{9a}$$

where $c_i'$ is the mutated i-th gene, $c_i$ is the original value of the i-th gene, Δ is the mutation range (set to 1.0 by default), and $\delta_i$ is the mutation factor determined by:

$$\delta_i = \begin{cases} 2u_i^{\frac{1}{\eta+1}} - 1 & \text{if } u_i \leq 0.5 \\ 1 - (2(1-u_i))^{\frac{1}{\eta+1}} & \text{if } u_i \geq 0.5 \end{cases}$$

where $u_i$ is a uniform random number in [0,1], and η (eta) is the distribution index (set to 2 by default). Each gene has a probability pm = 0.75 of being selected for mutation.

This mutation scheme introduces variability in a controlled manner, with the distribution index η determining the likelihood of creating solutions near or far from the parent. Lower values of η allow for more distant mutations, while higher values tend to create offspring closer to their parents. The scheme allows for both exploration and fine-tuning of the solution space, maintaining genetic diversity and preventing premature convergence.

***Crossover***

Crossover was performed at a rate of 75% using the Simulated Binary Crossover (SBX) mechanism on all sets of parents. Parents are adjacent individuals stored in the population matrix. SBX creates two offspring that are centered around their two parents in the decision space, with a spread proportional to the distance between the parents. The crossover operation can be mathematically represented as $c_1 = \mu - \beta\, c$ and ($c_2 = \mu + \beta\, c$), where ($\mu = \frac{v_1+v_2}{2}$)and ($c = \frac{v_1-v_2}{2}$) denote the mean and half the difference of the parent vectors $v_1$ and $v_2$, respectively. The spread factor β is defined as $\beta = (2\,\mathrm{URN})^{\frac{1}{\gamma+1}}$ when $\mathrm{URN} \leq 0.5$ or $\beta = \left(2\,(1-\mathrm{URN})\right)^{-\frac{1}{\gamma+1}}$ when $\mathrm{URN} > 0.5$, where URN is a uniform random number in the interval [0, 1] and γ is the distribution index (set to 2 by default). Additionally, each component has a recombination probability $p_m = 0.3$.This crossover scheme facilitates both exploration and exploitation of the search space; the distribution index γ controls how similar the offspring are to their parents—a larger γ results in offspring that are closer to the parents, while a smaller γ permits more diverse offspring. This mechanism is particularly effective for real-valued optimization as it preserves the average of the parent values while allowing for controlled variation in the offspring.

***Selection***

For the selection process, tournament selection was utilized. This method involves randomly selecting multiple subsets of individuals from the population and then choosing the fittest individual from each to be a part of the next generation. The primary advantage of tournament selection is its balance between maintaining diversity within the population and ensuring the propagation of the fittest individuals, and is less susceptible to premature convergence compared to other selection strategies [1]. We chose a tournament size of 5% of the overall population, dividing the total population of 100,000 individuals into 20 groups of 5000 individuals every generation.

Advantage over alternative methods:

GAs are renowned for their ability to effectively navigate and search through high-dimensional and non-linear parameter spaces, characteristics inherent to our biochemical oscillator model REF Dorsey and Mayer, Genetic algorithms 1995.

**Analytical Model reduction via Quasi Steady-state Approximations**

While the QSS reduction provides valuable analytical confirmation of necessary conditions, its limitations should be acknowledged. The derived algebraic expressions for free A and P can yield non-physical negative concentrations under certain initial total concentration regimes (specifically if Atot < Ltot or Ptot < Ltot), indicating boundaries where the QSS assumptions may break down stoichiometrically. Nonetheless, within its domain of validity, the mathematical analysis corroborates the conclusions from simulation studies: oscillations in this system fundamentally require both the rate enhancement conferred by membrane localization (DF > 1) and the complete feedback topology structured by the adaptor protein A, including the recruitment of both kinase and phosphatase. Here is a detailed breakdown of the QSS approach. First, we define to the rate of production of $x$ in reaction $\mathrm{rxn}$ as $\mathrm{Rate}_{x,n}$. Namely, summing over this rate in the whole system yields the time derivative of $x$:

$$\frac{dx}{dt} = \sum_{n=1}^{k} \mathrm{Rate}_{x,n}$$

Consider Michaelis-Menten reactions showing below within some big system of reactions, where $x$, $y$, $p$ are concentration of reactant $X$, $Y$ and $P$, and $i$ is the concentration of intermediate $I$ respectively.

$$X + Y \xleftrightarrow{k_1, k_{-1}} I \xrightarrow{k_2} P$$

Apply the QSS above procedurally following the protocol below:

| **Step 1**<br>LK in QSS;<br>LpAKL in QSS | **Step 2**<br>LpP in QSS;<br>LpAPL in QSS | **Step 3**<br>K in QSS;<br>LpA in QSS | **Final result** |
|---|---|---|---|
| (kfLK,krLK), L + K <--> LK<br>kcatLK, LK --> Lp + K<br>(kfLA,krLA), Lp + A <--> LpA | keff1, L + K -> Lp + K<br>(kfLA,krLA), Lp + A <--> LpA<br>(kfAK,krAK), LpA + K <--> LpAK | keff1, L + K -> Lp + K<br>(kfLA,krLA), Lp + A <--> LpA<br>*(kfAK,krAK), LpA + K <--> LpAK* | keff1, L + K -> Lp + K<br>(kfLA,krLA), Lp + A <--> LpA<br>*(kfAK,krAK), LpA + K <--> LpAK (implicit)* |

| | | | |
|---|---|---|---|
| (kfAK,krAK), LpA + K <--> LpAK<br>(k*y,krLK), LpAK + L <--> LpAKL<br>kcatLK, LpAKL --> Lp + LpAK<br>(kfLP,krLP), Lp + P <--> LpP<br>kcatLP, LpP --> L + P<br>(kfAP,krAP), LpA + P <--> LpAP<br>(kfLP*y,krLP), Lp + LpAP <--> LpAPLp<br>kcatLP, LpAPLp --> L + LpAP | keff1*y, LpAK + L --> Lp + LpAK<br>(kfLP,krLP), Lp + P <--> LpP<br>kcatLP, LpP --> L + P<br>(kfAP,krAP), LpA + P <--> LpAP<br>(kfLP*y,krLP), Lp + LpAP <--> LpAPLp<br>kcatLP, LpAPLp --> L + LpAP | (kfAP,krAP), LpA + P <--> LpAP<br>keff1*y, LpAK + L --> Lp + LpAK<br>keff7, Lp + P --> L + P<br>keff7*y, Lp + LpAP --> L + LpAP | (kfAP,krAP), LpA + P <--> LpAP<br>keff1*y, LpAK + L --> Lp + LpAK<br>keff7, Lp + P --> L + P<br>keff7*y, Lp + LpAP --> L + LpAP |
| $k_{\mathrm{eff1}} = \dfrac{k_f^{\mathrm{LK}} k_{\mathrm{cat}}^{\mathrm{LK}}}{k_r^{\mathrm{LK}} + k_{\mathrm{cat}}^{\mathrm{LK}}}$<br>$[\mathrm{LK}] = \dfrac{y k_{\mathrm{eff1}}}{k_{\mathrm{cat}}^{\mathrm{LK}}}[\mathrm{L}][\mathrm{LpAK}]$ | $k_{\mathrm{eff7}} = \dfrac{k_f^{\mathrm{LP}} k_{\mathrm{cat}}^{\mathrm{LP}}}{k_r^{\mathrm{LP}} + k_{\mathrm{cat}}^{\mathrm{LP}}}$<br>$[\mathrm{LpP}] = \dfrac{y k_{\mathrm{eff7}}}{k_{\mathrm{cat}}^{\mathrm{LP}}}[\mathrm{Lp}][\mathrm{LpAP}]$ | $[\mathrm{K}] = \dfrac{k_r^{\mathrm{AK}}[\mathrm{LpAK}]}{k_f^{\mathrm{AK}}[\mathrm{LpA}]}$<br>$[\mathrm{LpA}] = \dfrac{k_f^{\mathrm{LA}}[\mathrm{Lp}][\mathrm{A}] + k_r^{\mathrm{AP}}[\mathrm{LpAP}]}{k_r^{\mathrm{LA}} + k_f^{\mathrm{AP}}[\mathrm{P}]}$ | |

Table. 1. Procedure of model reduction applying QSS LK, LpAKL, LpP, LpAPL, K, and LpA; in the analytical solution $y$ refers to the dimensional factor.

The full reduced system of differential equations we get is:

$$
\begin{cases}
\dfrac{d[\mathrm{L}]}{dt} = -k_{\mathrm{eff1}}[\mathrm{L}][\mathrm{K}] - y k_{\mathrm{eff1}}[\mathrm{L}][\mathrm{LpAK}] + k_{\mathrm{eff7}}[\mathrm{Lp}][\mathrm{P}] + y k_{\mathrm{eff7}}[\mathrm{LpAP}][\mathrm{Lp}] \\
\dfrac{d[\mathrm{Lp}]}{dt} = k_{\mathrm{eff1}}[\mathrm{L}][\mathrm{K}] + y k_{\mathrm{eff1}}[\mathrm{L}][\mathrm{LpAK}] - k_{\mathrm{eff7}}[\mathrm{Lp}][\mathrm{P}] - y k_{\mathrm{eff7}}[\mathrm{LpAP}][\mathrm{Lp}] - k_f^{\mathrm{LA}}[\mathrm{Lp}][\mathrm{A}] + k_r^{\mathrm{LA}}[\mathrm{LpA}] \\
[\mathrm{P}]_{\mathrm{tot}} = [\mathrm{P}] + [\mathrm{LpAP}] + \dfrac{k_{\mathrm{eff7}}}{k_{\mathrm{cat}}^{\mathrm{LP}}}[\mathrm{Lp}][\mathrm{P}] + \dfrac{y k_{\mathrm{eff7}}}{k_{\mathrm{cat}}^{\mathrm{LP}}}[\mathrm{Lp}][\mathrm{LpAP}] \\
[\mathrm{K}]_{\mathrm{tot}} = \dfrac{k_{\mathrm{eff1}}}{k_{\mathrm{cat}}^{\mathrm{LK}}}[\mathrm{L}][\mathrm{K}] + \dfrac{y k_{\mathrm{eff1}}}{k_{\mathrm{cat}}^{\mathrm{LK}}}[\mathrm{L}][\mathrm{LpAK}] + [\mathrm{K}] + [\mathrm{LpAK}] \\
[\mathrm{A}]_{\mathrm{tot}} = [\mathrm{A}] + [\mathrm{LpA}] + [\mathrm{LpAP}] + [\mathrm{LpAK}] + \dfrac{y k_{\mathrm{eff7}}}{k_{\mathrm{cat}}^{\mathrm{LP}}}[\mathrm{Lp}][\mathrm{LpAP}] + \dfrac{y k_{\mathrm{eff1}}}{k_{\mathrm{cat}}^{\mathrm{LK}}}[\mathrm{L}][\mathrm{LpAK}] \\
[\mathrm{L}]_{\mathrm{tot}} = [\mathrm{L}] + [\mathrm{Lp}] + [\mathrm{LpA}] + [\mathrm{LpAP}] + [\mathrm{LpAK}] + \\
\qquad \dfrac{2 y k_{\mathrm{eff7}}}{k_{\mathrm{cat}}^{\mathrm{LP}}}[\mathrm{Lp}][\mathrm{LpAP}] + \dfrac{k_{\mathrm{eff7}}}{k_{\mathrm{cat}}^{\mathrm{LP}}}[\mathrm{Lp}][\mathrm{P}] + \dfrac{2 y k_{\mathrm{eff1}}}{k_{\mathrm{cat}}^{\mathrm{LK}}}[\mathrm{L}][\mathrm{LpAK}] + \dfrac{k_{\mathrm{eff1}}}{k_{\mathrm{cat}}^{\mathrm{LK}}}[\mathrm{L}][\mathrm{K}] \\
[\mathrm{K}] = \dfrac{k_r^{\mathrm{AK}}[\mathrm{LpAK}]}{k_f^{\mathrm{AK}}[\mathrm{LpA}]} \\
[\mathrm{LpA}] = \dfrac{k_f^{\mathrm{LA}}[\mathrm{Lp}][\mathrm{A}] + k_r^{\mathrm{AP}}[\mathrm{LpAP}]}{k_r^{\mathrm{LA}} + k_f^{\mathrm{AP}}[\mathrm{P}]}
\end{cases}
$$

We then do dimensional reduction for stability analysis. we set characteristic concentration to $[\mathrm{L}]_{\mathrm{tot}}$ and characteristic time to $(k_{\mathrm{eff1}}[\mathrm{L}]_{\mathrm{tot}})^{-1}$. All constants and variables can be converted to dimensionless via:

$$
\begin{cases}
l(\tau) = \dfrac{[\mathrm{L}]}{[\mathrm{L}]_{\mathrm{tot}}} \\
m(\tau) = \dfrac{[\mathrm{Lp}]}{[\mathrm{L}]_{\mathrm{tot}}} \\
\chi = \dfrac{[\mathrm{K}]}{[\mathrm{L}]_{\mathrm{tot}}} \\
a = \dfrac{[\mathrm{A}]}{[\mathrm{L}]_{\mathrm{tot}}} \\
p = \dfrac{[\mathrm{P}]}{[\mathrm{L}]_{\mathrm{tot}}} \\
X = \dfrac{[\mathrm{K}]_{\mathrm{tot}}}{[\mathrm{L}]_{\mathrm{tot}}} \\
A = \dfrac{[\mathrm{A}]_{\mathrm{tot}}}{[\mathrm{L}]_{\mathrm{tot}}} \\
P = \dfrac{[\mathrm{P}]_{\mathrm{tot}}}{[\mathrm{L}]_{\mathrm{tot}}} \\
\tau = k_{\mathrm{eff1}}[\mathrm{L}]_{\mathrm{tot}}t \\
\kappa_1 = \dfrac{k_{\mathrm{eff1}}}{k_{\mathrm{eff1}}} = 1 \\
\kappa_7 = \dfrac{k_{\mathrm{eff7}}}{k_{\mathrm{eff1}}} \\
\kappa_{a2} = \dfrac{k_f^{\mathrm{LA}}}{k_{\mathrm{eff1}}} \\
\kappa_{a4} = \dfrac{k_f^{\mathrm{AP}}}{k_{\mathrm{eff1}}} \\
\kappa_{b2} = \dfrac{k_r^{\mathrm{LA}}}{k_{\mathrm{eff1}}[\mathrm{L}]_{\mathrm{tot}}} \\
\kappa_{b4} = \dfrac{k_r^{\mathrm{AP}}}{k_{\mathrm{eff1}}[\mathrm{L}]_{\mathrm{tot}}} \\
K_3 = \dfrac{k_f^{\mathrm{AK}}}{k_r^{\mathrm{AK}}}[\mathrm{L}]_{\mathrm{tot}} \\
\dfrac{1}{\kappa_{c1}} = \dfrac{k_{\mathrm{cat}}^{\mathrm{LK}}\kappa_1}{k_{\mathrm{eff1}}[\mathrm{L}]_{\mathrm{tot}}} \\
\dfrac{1}{\kappa_{c7}} = \dfrac{k_{\mathrm{cat}}^{\mathrm{LP}}\kappa_7}{k_{\mathrm{eff1}}[\mathrm{L}]_{\mathrm{tot}}}
\end{cases}
$$

Theta denotes dimensionless concentration for all other variables. The range of the variables are: $l, m \in [0,1]$ and $l + m \in [0, 1]$, $\tau, \kappa_7, \kappa_{ai}, \kappa_{bi} \in [0, \infty)$. Therefore, the concentrations can be expressed as dimensionless expressions times the total concentration of species containing L:

$$\begin{cases} \dfrac{dl}{d\tau} = -l\chi - yl\theta_{\text{LpAK}} + \kappa_7 mp + y\kappa_7 m\theta_{\text{LpAP}} \\ \dfrac{dm}{d\tau} = l\chi + yl\theta_{\text{LpAK}} - \kappa_7 mp - y\kappa_7 m\theta_{\text{LpAP}} - \kappa_{a2}ma + \kappa_{b2}\theta_{\text{LpA}} \\ P = p + \theta_{\text{LpAP}} + \kappa_{c7}mp + y\kappa_{c7}m\theta_{\text{LpAP}} \\ X = \kappa_{c1}l\chi + y\kappa_{c1}l\theta_{\text{LpAK}} + \chi + \theta_{\text{LpAK}} \\ A = a + \theta_{\text{LpA}} + \theta_{\text{LpAP}} + \theta_{\text{LpAK}} + y\kappa_{c7}m\theta_{\text{LpAP}} + y\kappa_{c1}l\theta_{\text{LpAK}} \\ 1 = l + m + \theta_{\text{LpA}} + \theta_{\text{LpAP}} + \theta_{\text{LpAK}} \\ \quad +2y\kappa_{c7}m\theta_{\text{LpAP}} + 2y\kappa_{c1}l\theta_{\text{LpAK}} + \kappa_{c7}mp + \kappa_{c1}l\chi \\ \chi = \dfrac{\theta_{\text{LpAK}}}{K_3\theta_{\text{LpA}}} \\ \theta_{\text{LpA}} = \dfrac{\kappa_{a2}ma + \kappa_{b4}\theta_{\text{LpAP}}}{\kappa_{b2} + \kappa_{a4}p} \end{cases}$$

Note this is the 2D reduced system. The system $(l, m)\sim\tau$ is 2D autonomous. However, the analytical solution to this full system via Mathematica yields a system of differential equations that is gigabytes large, which is far too complicated for any further assumption. Here we make the following simplification: replacing mass conservation of P, K, A with strict constant P, constant K, and constant A and QSS of K with constant LpA.

$$\begin{cases} \dfrac{dl}{d\tau} = -l\chi - yl\theta_{\text{LpAK}} + \kappa_7 mp + y\kappa_7 m\theta_{\text{LpAP}} \\ \dfrac{dm}{d\tau} = l\chi + yl\theta_{\text{LpAK}} - \kappa_7 mp - y\kappa_7 m\theta_{\text{LpAP}} - \kappa_{a2}ma + \kappa_{b2}\theta_{\text{LpA}} \\ P = p \\ X = \chi \\ A = a \\ 1 = l + m + \theta_{\text{LpA}} + \theta_{\text{LpAP}} + \theta_{\text{LpAK}} \\ \quad +2y\kappa_{c7}m\theta_{\text{LpAP}} + 2y\kappa_{c1}l\theta_{\text{LpAK}} + \kappa_{c7}mp + \kappa_{c1}l\chi \\ \theta_{\text{LpA0}} = \theta_{\text{LpA}} \\ \theta_{\text{LpA}} = \dfrac{\kappa_{a2}ma + \kappa_{b4}\theta_{\text{LpAP}}}{\kappa_{b2} + \kappa_{a4}p} \end{cases}$$

Now we can write the further simplified 2D autonomous system as:

$$\begin{cases} \dfrac{dl}{d\tau} = \dfrac{1}{\kappa_{b4} + 2y\kappa_{b4}\kappa_{c1}l}\big(y\kappa_{b4}(1 - X\kappa_{c1})l^2 + \kappa_7 m(y\theta_{\text{LpA0}}(P\kappa_{a4} + \kappa_{b2}) + P\kappa_{b4} \\ \quad -Ay\kappa_{a2}m) + l(y\theta_{\text{LpA0}}(P\kappa_{a4} + \kappa_{b2}) - (X + y - y\theta_{\text{LpA0}})\kappa_{b4} \\ \quad +ym(-A\kappa_{a2} + \kappa_{b4} + 2y\theta_{\text{LpA0}}(P\kappa_{a4} + \kappa_{b2})(\kappa_7\kappa_{c1} + \kappa_{c7}) \\ \quad +P\kappa_{b4}(2\kappa_7\kappa_{c1} + \kappa_{c7}) - 2Ay\kappa_{a2}(\kappa_7\kappa_{c1} + \kappa_{c7})m))\big) \\ \dfrac{dm}{d\tau} = \theta_{\text{LpA0}}\kappa_{b2} + Xl - P\kappa_7 m - A\kappa_{a2}m + \dfrac{y\kappa_7 m\big(-\theta_{\text{LpA0}}(P\kappa_{a4} + \kappa_{b2}) + A\kappa_{a2}m\big)}{\kappa_{b4}} \\ \quad -\dfrac{1}{\kappa_{b4} + 2y\kappa_{b4}\kappa_{c1}l}yl(-\kappa_{b4} + \theta_{\text{LpA0}}(P\kappa_{a4} + \kappa_{b2} + \kappa_{b4}) + (\kappa_{b4} + X\kappa_{b4}\kappa_{c1})l + \\ \quad m(-A\kappa_{a2} + \kappa_{b4} + 2y\theta_{\text{LpA0}}(P\kappa_{a4} + \kappa_{b2})\kappa_{c7} + P\kappa_{b4}\kappa_{c7} - 2Ay\kappa_{a2}\kappa_{c7}m)) \end{cases}$$

**Existence of limit cycle in further simplified 2D system for some parameter combination**

Using system above, we prove the existence of limit cycle for some parameter combination by simply giving an example. Consider the following set of parameters:

| | | |
|---|---|---|
| X = 0.321551891; | $\kappa_{a2}$ = 535.2958859; | $K_3$ = 0.054087196; |
| P = 0.112776887; | $\kappa_{b2}$ = 55.65925966; | $\kappa_{c1}$ = 0.000441364; |
| A = 1; | $\kappa_{a4}$ = 3.351166619; | $\kappa_{c7}$ = 0.559693066; |
| y = 1000.0; | $\kappa_{b4}$ = 90.20379571; | $\kappa_7$ = 269.2319406; |
| | | $\theta_{LpA0}$ = 0.05; |

**Effect of changes on the system on limit cycle in further simplified 2D system**

Here we verify analytically that some changes on the system will lead to absence of limit cycle the whole regime. First, reminder of a few corollaries that we are using in this section:

**Corollary 1: No oscillation in simple autonomous system**

For any 2D autonomous system $\frac{dx}{dt} = f(x, y), \frac{dy}{dt} = g(x, y)$ , if $\frac{g(x,y)}{f(x,y)} = c$, where $c$ is a real constant, then there is no limit cycle in the system.

**Index Theory (Strogatz 6.8)**

Any closed orbit in the phase plane must enclose fixed points whose indices sum to +1.

Corollary 2.1:
If there is no fixed point in the domain of interest and not on the boundary (l in (0,1) and m in (0,1)), there is no limit cycle in the domain of interest.

Corollary 2.2:
If there exists only 1 fixed point in the domain of interest and not on the boundary (l in (0,1) and m in (0,1)), and the determinant of the Jacobian evaluated at this fixed point is smaller than zero, then there is no limit cycle in the domain of interest.

**Bendixson–Dulac theorem (Burton, Theodore Allen p.318)**

If there exists a Dulac function $\phi(x, y)$ such that the expression $\frac{\partial(\phi f)}{\partial x} + \frac{\partial(\phi g)}{\partial y}$ has the same sign ($\neq 0$) everywhere except possibly in a set of measure 0 in a simply connected region of the plane, then the plane autonomous system $\frac{dx}{dt} = f(x, y), \frac{dy}{dt} = g(x, y)$ has no nonconstant periodic solutions lying entirely within the region.

Corollary 3:
If trace of Jacobian evaluated everywhere in the domain of interest has the same sign, then there is no limit cycle in the domain of interest. (Plug in $\phi(x, y) = 1$)

**Poincaré–Andronov–Hopf bifurcation**

A local bifurcation in which a fixed point of a dynamical system loses stability, as a pair of complex conjugate eigenvalues—of the linearization around the fixed point—crosses the complex plane imaginary axis as a parameter crosses a threshold value.

Corollary 4:

In 2D plane autonomous system, if this local bifurcation exists, then for some parameter combination there exists a limit cycle.

**Case 1: delete A**

If A is deleted from the system, then both the concentration of A and LpA are equal to zero. Therefore, in the dimensionless system, $A = 0$ and $\theta_{\mathrm{LPA0}} = 0$.

$$\begin{cases} \dfrac{dl}{d\tau} = -l\chi - yl\theta_{\mathrm{LpAK}} + \kappa_7 mp + y\kappa_7 m\theta_{\mathrm{LpAP}} \\ \dfrac{dm}{d\tau} = l\chi + yl\theta_{\mathrm{LpAK}} - \kappa_7 mp - y\kappa_7 m\theta_{\mathrm{LpAP}} - \kappa_{a2} ma + \kappa_{b2}\theta_{\mathrm{LpA}} \\ P = p \\ X = \chi \\ 0 = a \\ 1 = l + m + \theta_{\mathrm{LpA}} + \theta_{\mathrm{LpAP}} + \theta_{\mathrm{LpAK}} \\ \quad +2y\kappa_{c7} m\theta_{\mathrm{LpAP}} + 2y\kappa_{c1} l\theta_{\mathrm{LpAK}} + \kappa_{c7} mp + \kappa_{c1} l\chi \\ 0 = \theta_{\mathrm{LpA}} \\ \theta_{\mathrm{LpA}} = \dfrac{\kappa_{a2} ma + \kappa_{b4}\theta_{\mathrm{LpAP}}}{\kappa_{b2} + \kappa_{a4} p} \end{cases}$$

Therefore, the system of equation simplifies to follows:

$$\begin{cases} \dfrac{dl}{d\tau} = -Xl + P\kappa_7 m - \dfrac{yl(1 - l - X\kappa_{c1} l - m - P\kappa_{c7} m)}{1 + 2y\kappa_{c1} l} \\ \dfrac{dm}{d\tau} = Xl - P\kappa_7 m + \dfrac{yl(1 - l - X\kappa_{c1} l - m - P\kappa_{c7} m)}{1 + 2y\kappa_{c1} l} \end{cases}$$

Note that here $\frac{dl}{d\tau} = -\frac{dm}{d\tau}$, this means that $\frac{dl}{dm} = -1$, which is a real constant. By Corollary 1, there is no limit cycle in the system.

**Case 2: delete P**

If P is deleted from the system, then both the concentration of P and LpAP are set to zero. Therefore, $\theta_{\mathrm{LpAP}} = 0$ and $P = 0$. However, this results in $\theta_{\mathrm{LpA0}} = \frac{\kappa_{a2} mA}{\kappa_{b2}}$ which means $m(\tau)$ becomes a constant. (?)

**Case 3: set DF=1**

Short answer: Empirically from numerical calculations, we do not observe limit cycles. However analytically, with dimensionality factor (y=1) it is hard to decisively conclude whether this system allows a limit cycle, according to Bendixson–Dulac theorem.

Write the system as $\dot{l} = F_1(l, m), ; \dot{m} = F_2(l, m)$ and a Dulac function to use is:

$$D(l) = \kappa_{b4}(1 + 2\kappa_{c1}l) \qquad \left(> 0 \text{ for } l > -\frac{1}{2\kappa_{c1}}\right)$$

Now based on the form of equation let:

$$F_1 = (-Xl + P\kappa_7 m) + \Phi(l,m) = F_{1A} + \frac{1}{D}\Phi(l,m)$$

Where $F_{1A}$ denotes the “affine part” of $F_1$ which is $-Xl + P\kappa_7 m$ and $\Phi(l,m)/D$ denotes the nonlinear part. Similarly let:

$$F_2 = \left(\theta_{\text{LpA0}}\kappa_{b2} + Xl - P\kappa_7 m - A\kappa_{a2}m + \frac{\kappa_7 m}{\kappa_{b4}}\left(-\theta_{\text{LpA0}}(P\kappa_{a4} + \kappa_{b2}) + A_{\kappa 2}m\right)\right) - \frac{l}{D}\Psi(l,m)$$
$$= F_{2AQ} - \frac{l}{D}\Psi(l,m)$$

Where $F_{1A}$ denotes the “affine and quadratic part” of $F_2$ and $-l\Psi(l,m)/D$ denotes the nonlinear part. Note that through a simple but lengthy polynomial check we can get:

$$\frac{\partial \Phi}{\partial l} = \frac{\partial (l\Psi)}{\partial m}$$

Compute the Dulac divergence:

$$\nabla \cdot (DF) = \frac{\partial}{\partial l}(DF_1) + \frac{\partial}{\partial m}(DF_2) = \frac{\partial}{\partial l}\big(DF_{1A} + \Phi(l,m)\big) + \frac{\partial}{\partial m}\big(DF_{2AQ} - \Psi(l,m)\big)$$

Simplify, we get:

$$\nabla \cdot (DF) = \left(\frac{\partial \Phi}{\partial l} - \frac{\partial (l\Psi)}{\partial m}\right) + \left(\frac{\partial}{\partial l}(DF_{1A}) + \frac{\partial}{\partial m}(DF_{2AQ})\right) = \frac{\partial}{\partial l}(DF_{1A}) + \frac{\partial}{\partial m}(DF_{2AQ})$$

Expanding the derivatives, we can prove that the sign of $\nabla \cdot (DF)$ does not always stay positive or negative and thus the result is indecisive. However, it is only one case of a specific Dulac function, and we acknowledge that there might exist a Dulac function that can prove the absence of limit cycle.

**Case 4: set $\boldsymbol{\kappa_{b2}}$=0**

The system becomes:

$$\begin{cases} \dfrac{dl}{d\tau} = -l\chi - yl\theta_{\mathrm{LpAK}} + \kappa_7 mp + y\kappa_7 m\theta_{\mathrm{LpAP}} \\ \dfrac{dm}{d\tau} = l\chi + yl\theta_{\mathrm{LpAK}} - \kappa_7 mp - y\kappa_7 m\theta_{\mathrm{LpAP}} - \kappa_{a2} ma \\ P = p \\ X = \chi \\ A = a \\ 1 = l + m + \theta_{\mathrm{LpA}} + \theta_{\mathrm{LpAP}} + \theta_{\mathrm{LpAK}} \\ +2y\kappa_{c7} m\theta_{\mathrm{LpAP}} + 2y\kappa_{c1} l\theta_{\mathrm{LpAK}} + \kappa_{c7} mp + \kappa_{c1} l\chi \\ \theta_{\mathrm{LpA0}} = \theta_{\mathrm{LpA}} \\ \theta_{\mathrm{LpA}} = \dfrac{\kappa_{a2} ma + \kappa_{b4}\theta_{\mathrm{LpAP}}}{\kappa_{b2} + \kappa_{a4} p} \end{cases}$$

Analytical methods do not rule out limit cycle and numerical solution shows evidence for existence of limit cycle, but this result might be a numerical artefact due to its instability:

| | | |
|---|---|---|
| X = 0.321551891; | $\kappa_{a2}$ = 535.2958859; | $K_3$ = 0.054087196; |
| P = 0.112776887; | $\kappa_{b2}$ = 0.0; | $\kappa_{c1}$ = 0.000441364; |
| A = 1; | $\kappa_{a4}$ = 3.351166619; | $\kappa_{c7}$ = 0.559693066; |
| y = 1000.0; | $\kappa_{b4}$ = 90.20379571; | $\kappa_7$ = 269.2319406; |
| | | $\theta_{LpA0}$ = 0.05; |

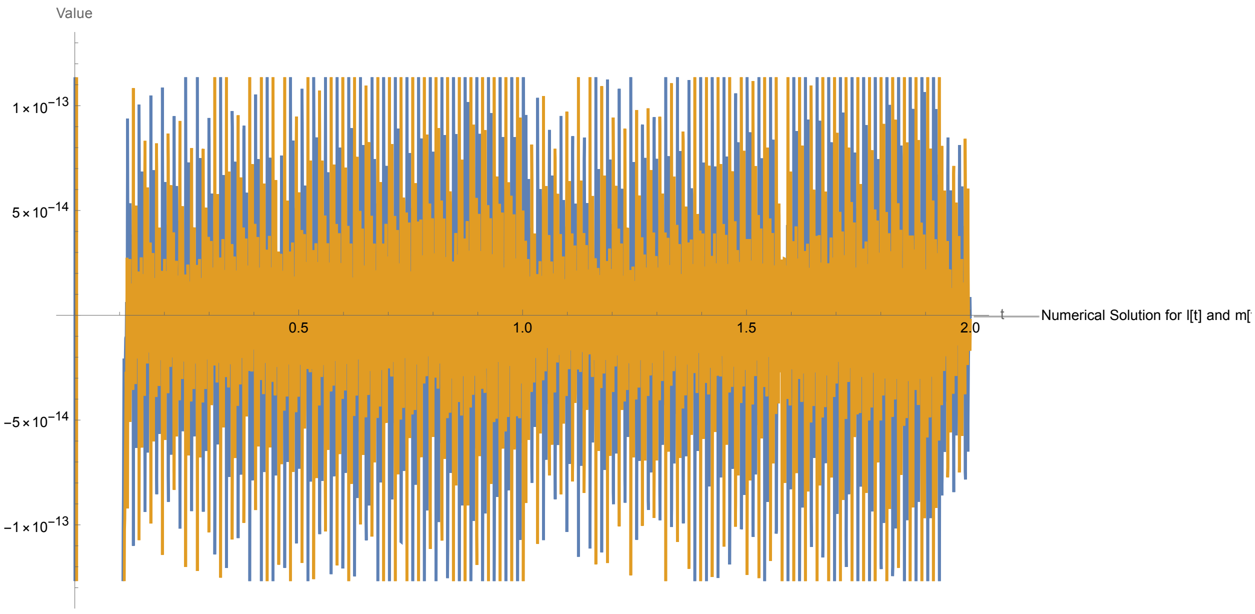


**Case 5: set $\kappa_{a4}$=0, $\theta_{\mathrm{LpAP}} = 0$**

Trivial linear case. No limit cycle in the system because $m = \frac{\theta_{\mathrm{LpA0}}\kappa_{b2}}{A\kappa_{a2}}$ is constant under this assumption. The full system is:

$$
\begin{cases}
\frac{dl}{d\tau} = -l\chi - yl\theta_{\text{LpAK}} + \kappa_7 mp \\
\frac{dm}{d\tau} = l\chi + yl\theta_{\text{LpAK}} - \kappa_7 mp - \kappa_{a2} ma + \kappa_{b2}\theta_{\text{LpA}} \\
P = p \\
X = \chi \\
A = a \\
1 = l + m + \theta_{\text{LpA}} + \theta_{\text{LpAK}} \\
+2y\kappa_{c7} m\theta_{\text{LpAP}} + 2y\kappa_{c1} l\theta_{\text{LpAK}} + \kappa_{c7} mp + \kappa_{c1} l\chi \\
\theta_{\text{LpA0}} = \theta_{\text{LpA}} \\
\theta_{\text{LpA}} = \frac{\kappa_{a2} ma}{\kappa_{b2}}
\end{cases}
$$

**Case 6: set $\boldsymbol{\kappa_{a2}}$=0, $\boldsymbol{\theta_{\text{LpA}} = 0}$**

Trivial linear case. No limit cycle in the system because $\frac{\frac{dl}{d\tau}}{\frac{dm}{d\tau}} = -1$ which is constant under this assumption (corollary 1). The full system is:

$$
\begin{cases}
\frac{dl}{d\tau} = -l\chi - yl\theta_{\text{LpAK}} + \kappa_7 mp + y\kappa_7 m\theta_{\text{LpAP}} \\
\frac{dm}{d\tau} = l\chi + yl\theta_{\text{LpAK}} - \kappa_7 mp - y\kappa_7 m\theta_{\text{LpAP}} \\
P = p \\
X = \chi \\
A = a \\
1 = l + m + \theta_{\text{LpA}} + \theta_{\text{LpAP}} + \theta_{\text{LpAK}} \\
+2y\kappa_{c7} m\theta_{\text{LpAP}} + 2y\kappa_{c1} l\theta_{\text{LpAK}} + \kappa_{c7} mp + \kappa_{c1} l\chi \\
\theta_{\text{LpA0}} = 0 \\
0 = \frac{\kappa_{b4}\theta_{\text{LpAP}}}{\kappa_{b2} + \kappa_{a4} p}
\end{cases}
$$

**Stability analysis using the Robust Lyapunov Function approach[2,3]**

This approach supports that the full post-translational modification cycle (the system without A) reaches a global steady-state, and thus does not oscillate. However, with the addition of A that binds the membrane and either of the enzymes, no stability can be established using this general approach.

**Experimental Data from Ezra:**

| Binding of PIP2 to AP2 (Höning et al., 2005) | Binding of AP2 Subunit to PIP5K (Krauss et al., 2006) | Binding of Synaptojanin 1 to AP2 α-Appendage (Praefcke et al., 2004) | Catalytic Activity of Synaptojanin 1 on PIP2 (Paesmans et al., 2020) | Catalytic Activity of PIP5K1C (Shulga et al., 2012) |
|---|---|---|---|---|
| $k_{on}$= 0.7 × $10^{-3}$ μ$M^{-1}s^{-1}$ | $k_{on}$= 0.118 μ$M^{-1}s^{-1}$ | $K_d$= 29 μM | $K_M$ = 39 μM | $K_M$ = 15 μM |
| $k_{off}$= 2.0 × $10^{-3}$ $s^{-1}$ | $k_{off}$= 0.609 $s^{-1}$ | | $k_{cat}$ = 85.3 $s^{-1}$ | |

**Table 1: Experimental Values.** In simulations, the value of $k_{cat}$ for PIP5K was determined using the formula $k_{cat} = ( K_M \times k_{on} ) - k_{off}$ , where $k_{on}$ and $k_{off}$ were sampled from a range of physically realistic values that did not lead to a negative catalytic rate constant. Similarly, $k_{on}$ for Synaptojanin 1 was sampled from physically realistic ranges, and $k_{off}$ was calculated via $k_{off} = ( K_M \times k_{on} ) - k_{cat}$. $k_{on}$ for the binding of Synaptojanin to AP2 was also sampled from physically realistic ranges, and $k_{off}$ was calculated using $k_{off} = k_{on} \times K_d$. McIntire et al. report different values for the catalytic activity of Synaptojanin 1 on $PIP_2$ ($K_M$ = 138.5 μM, $k_{cat}$= 50. $s^{-1}$), though these were not used in our investigation (2013).

*Note koff for the AP2 binding to PIP5K should be 0.0609 s-1, or 10x slower.

### Results from optimizing individuals under experimental constraints

Results were only found for the highest values of DF. The above values were all constrained, and all other variables were free.

The oscillations in Amem were typically quite minimal, despite visible oscillations in the lipid species.

The solution below is at the border of the numerically sampled accessible region in Fig 3.

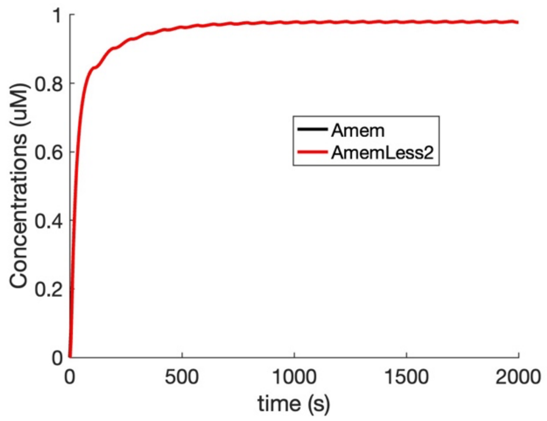


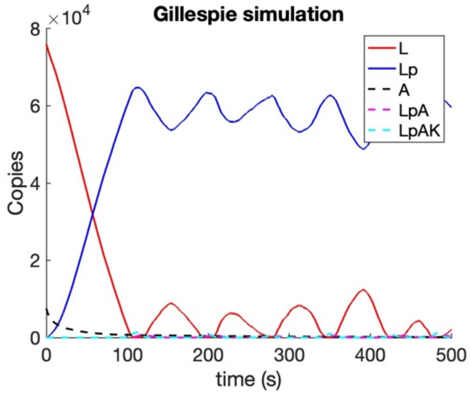


**NERDSS model molecule geometry and diffusion constants**

For all bulk solution species, D = 25μm$^2$/s, Dr = 0.5rad$^2$/μs. For the lipid, D = 1 μm$^2$/s.

**Adaptor:**

```
COM     0.0000    0.0000    0.0000
lbs     0.0000    0.0000   -2.0000
ebs     0.0000    1.0000   -1.0000
```

**Kinase:**

```
COM     0.0000    0.0000    0.0000
abs     0.0000    0.0000    0.5000
lbs     0.0000    0.0000   -0.5000
```

**Phosphatase:**

| | | | |
|---|---|---|---|
| COM | 0.0000 | 0.0000 | 0.0000 |
| abs | 0.0000 | 0.0000 | 0.5000 |
| lbs | 0.0000 | 0.0000 | -0.5000 |

**Lipid:**

| | | | |
|---|---|---|---|
| COM | 0.0000 | 0.0000 | 0.0000 |
| head | 0.0000 | 0.0000 | 1.0000 |

**SUPPLEMENTAL FIGURES**

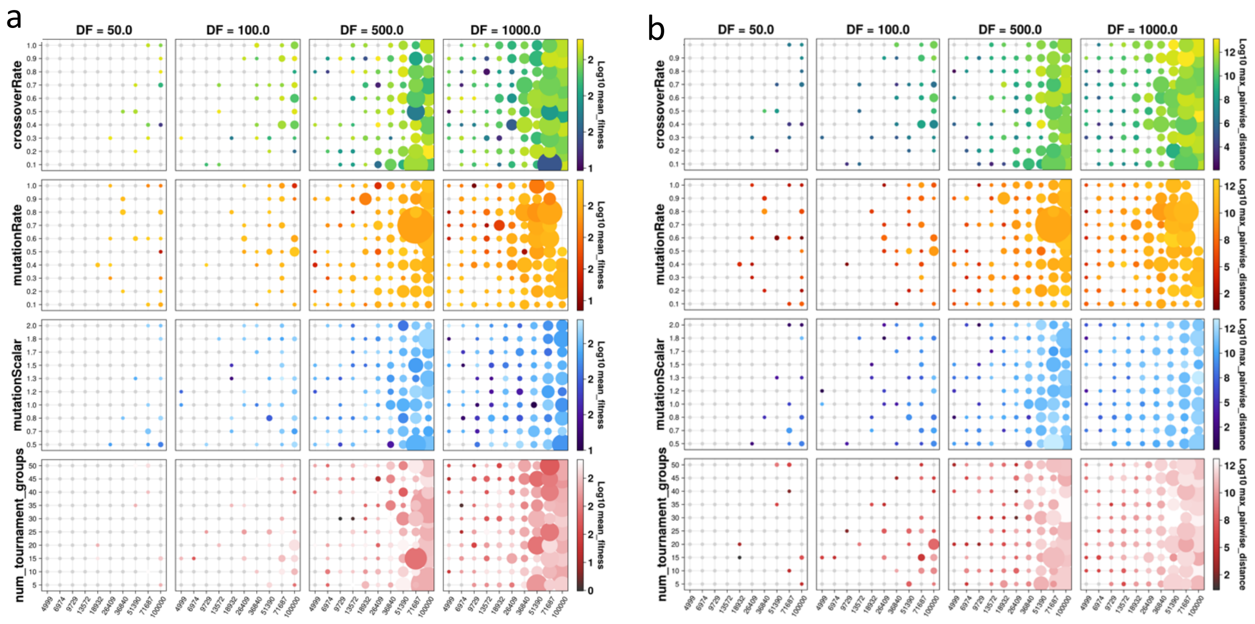


**SI Figure 1. Hyperparameter sweep indicates population size as the dominant factor for search quality and coverage.** Each row represents a hyperparameter that is varied as the others are held constant. Each column corresponds to hyperparameter sweep at different DF values. The x-axis of each grid is population size, while the y-axis is the row-specific hyperparameter value. Bubble size indicates absolute number of points found during optimization, while the colormaps indicate the performance metric for either quality or coverage. (a) Average fitness of the final populations for each of the hyperparameter grid points. Trends along the y-axes of each row of grids are unclear, though higher values for all the hyperparameters tested tend to show higher average fitness. (b) Log10 of the maximum pairwise distance in the final populations for each grid point. Maximum pairwise distance is even more homogenous than average fitness along the y-axis, and together with the average fitness grid plots show population size along the x-axes as the dominant factor in both performance metrics.

**TABLE**

| **Hyperparameter (units)** | **value** |
|---|---|
| Mutation rate per individual | 0.95 |
| Mutation probability (pm) per gene in an individual | 0.75 |
| Sampling parameter: Mutation distribution using Polynomial Mutation Scheme with parameter $\eta$ | 2 |
| Sampling parameter: Mutation range (delta) | 1.0 |
| Crossover rate for an individual (in addition to mutation) | 0.75 |
| SBX probability (pm): probability of choosing a gene in an individual | 0.3 |
| SBX distribution index ($\gamma$) | 2 |
| Tournament group size (# of individuals) | 5,000 |

| | |
|---|---|
| Population size (# of individuals) | 100,000 |

| Rate constants | Range |
|---|---|
| $\boldsymbol{k}_{\mathbf{f}}$ (binding rate) | 0.001 – 10 µ$M^{-1}s^{-1}$ |
| $\boldsymbol{k}_{\mathbf{r}}$(unbinding rate) | 0.001 – 1000 $s^{-1}$ |
| $\boldsymbol{k}_{\mathbf{cat}}$ (catalytic rate) | 0.001 – 1000 $s^{-1}$ |
| **Species** | **Range** |
| L/Lp | 0.1 – 100.0µ$M$ |
| K | 0.001 – 100.0µ$M$ |
| P | 0.001 – 100.0µ$M$ |
| A | 0.001 – 100.0µ$M$ |
